# Molecular mechanisms of stress-induced reactivation in mumps virus condensates

**DOI:** 10.1101/2021.07.10.451879

**Authors:** Xiaojie Zhang, Sindhuja Sridharan, Ievgeniia Zagoriy, Christina Eugster Oegema, Cyan Ching, Tim Pflaesterer, Herman K. H. Fung, Ina Poser, Christoph W. Müller, Anthony A. Hyman, Mikhail M. Savitski, Julia Mahamid

## Abstract

Many viruses can establish long-term persistent infections in the human host and cause chronic diseases. Here we combine cell biology, whole-cell proteomics and cryo-electron tomography to uncover how cellular stress disrupts the host-virus equilibrium in persistent infection and induces viral replication, using a model of negative-stranded RNA viruses, the mumps virus. We show that phosphorylation of the largely disordered viral phosphoprotein coincides with increased partitioning of viral polymerase into pre-formed liquid-like condensates and the formation of a stable replication machinery. By obtaining the first atomic models for the authentic mumps virus nucleocapsid, we elucidate a concomitant conformational change that exposes the viral genome to its replication machinery. These events that occur within viral condensates upon stress, together with concerted down-regulation of the host antiviral defense, provide an environment that supports up-regulation of viral replication and constitute a stress-mediated switch that disrupts the host-virus equilibrium in persistent infection.

**In Brief:** A multi-scale approach uncovers molecular and structural basis of how cellular stress provokes activation of persistent viral infection mediated by biomolecular condensates.

## Introduction

Long-term persistent viral infections are characterized by a metastable balance between viral replication and the host defense (Virgin et al., 2009). Stress to the host has been implicated in tipping this balance in a few cases, including the DNA viruses herpes simplex and human immunodeficiency viruses, and the emergence of pathology (Padgett et al., 1998; Piette and Legrand-Poels, 1994). However, the molecular mechanisms that underlie viral reactivation pertaining to stress remain elusive. An accidental encounter of a persistent viral infection in our cell cultures used for a study on stress response allowed us to investigate the cellular, molecular and structural basis of stress-mediated reactivation in a member of the negative-stranded RNA virus family *Paramyxoviridae*, the mumps virus (MuV).

Mumps is a highly contagious viral illness that was once a common childhood disease. Following implementation of widespread vaccination since the late 1960s, global incidence of mumps decreased substantially, and the disease was on the verge of elimination by 2001 (Rubin et al., 2015). Over the past decade, however, a rise in mumps cases became alarming because of vaccine hesitancy (Rubin et al., 2015; Su et al., 2020). Mumps virus (MuV) is self-limited; the majority of patients experience full recovery and obtain lifelong immunity (Su et al., 2020). However, chronic complications such as myositis (Chou, 1986), chronic arthritis (Huppertz and Chantler, 1991) and encephalitis (Julkunen et al., 1985) occur in MuV-infected patients, causing disabling long-term sequelae (dysfunction in muscle, joint tissue and the central nervous system) as consequences of persistent infection. These diseases do not arise due to an immediate effect of the infection, such as in parotitis, but develop months or years after natural infection, or vaccination in immunodeficient patients (Morfopoulou et al., 2017).

The MuV RNA genome encodes for the nucleocapsid (N), viral/phospho- (V/P, co-transcriptional products), matrix (M), fusion (F), small hydrophobic (SH), hemagglutinin-neuraminidase (HN) and large (L) proteins. N self-assembles into a helical capsid that accommodates the RNA genome to form a nucleocapsid, which protects the viral genome from host nucleases and serves as the template for viral replication. Both genome transcription and replication are carried out in cytoplasmic viral factories (Duc-Nguyen and Rosenblum, 1967) by the viral RNA-dependent RNA polymerase L, and require the phosphoprotein P that bridges between the nucleocapsid and the polymerase (Cox et al., 2014). The remaining MuV proteins are associated with nucleocapsid attachment to the inner side of the plasma membrane (HN), viral assembly (M), viral budding (F) and immunomodulation (V). Indeed, in cells persistently infected by the MuV or its family member measles virus (MeV), induction of interferon (IFN) signaling pathway effector protein as part of the host immune response is reported to be suppressed (Fujii et al., 1990). This is thought to be mediated by the V protein that directly disrupts the formation of the signal transducer and activator of transcription (STAT) 1-STAT2 complex (Rosas-Murrieta et al., 2010), or the STAT1, receptor-activated C kinase (RACK1) and IFN receptor complex (Kubota et al., 2002). On the other hand, viral replication in persistent MuV (Kristensson et al., 1984; Huppertz and Chantler, 1991) and MeV (Doi et al., 2016) infection in human joint tissue and mouse neurons has been shown to be self-restricting, suggesting a balance between the host immune response and viral replication at the chronic infection phase (Virgin et al., 2009).

Viral replication factories are subcellular microenvironments where viral genomes and replication machineries concentrate. The formation of viral factories is a complex process that has independently evolved in a variety of non-related viruses to increase the efficiency of viral replication (Etibor et al., 2021). Biomolecular condensation mediated by liquid-liquid phase separation of viral proteins with intrinsically disordered regions (IDRs) (Wang et al., 2018; Zhou et al., 2019) and viral genomic RNAs (Iserman et al., 2020; Zhou et al., 2019), has been recently shown to be fundamental for the organization of viral factories, regulation of viral replication and promotion of viral assembly in MeV (Zhou et al., 2019), rabies virus (Nikolic et al., 2017), vesicular stomatitis virus (Heinrich et al., 2018), and SARS-CoV-2 (Iserman et al., 2020; Lu et al., 2021). MuV replication factories are reported to be membrane-less enclusions as detected with conventional immuno-electron microscopy (EM) on chemically fixed sections of cells (Duc-Nguyen and Rosenblum, 1967). After viral inoculation, cytoplasmic MuV factories typically contain curved tubule-like structures, the nucleocapsids. However, biopsies from chronic myositis patient tissues show inclusions of MuV nucleocapsids, which appear to be more straight compared to their curved morphology during acute infection (Chou, 1986). Understanding the structural differences between these distinct forms of MuV nucleocapsids within the viral replication factories is important for gaining insights into their functional states and relevance to chronic diseases. However, such understanding has been limited due to the low resolution that immuno-EM can provide, as well as the lack of biological context when obtaining high-resolution cryo-EM structures with purified viral complexes (Shan et al., 2021).

Here, we combine light microscopy, whole-cell mass spectrometry (MS), cellular cryo-electron tomography (ET) and structural analysis, to provide an across-scales understanding of the cellular, molecular and structural changes in the virus and the host in persistent infection upon stress. We confirm that exogenous stress to the host leads to increased replication of the MuV in persistent infection. We report that intracellular MuV replication factories are liquid-like condensates that change their behavior and coarsen under stress. Using recent advances in cryo-ET (Qu et al., 2021; Tegunov et al., 2021) in combination with cryo-focused ion beam (FIB) milling (Mahamid et al., 2016; Wang et al., 2021), we visualize the molecular organization of viral factories in different cellular states, along with detailed structural insights into stress-related conformational changes in the MuV nucleocapsid. These findings suggested alterations in the protein interaction network within the viral condensates upon stress, which we dissected using a proteomics approach to reveal concerted changes in viral protein phosphorylation and solubility, that coincide with stabilization of the viral replication machinery. We thus demonstrate molecular and structural changes that link up-regulation of viral replication within cellular condensates to exogenous stress to the host.

## Results

### Viral replication and release are accelerated by stress

Persistent infection of *Paramyxoviridae* can be established within weeks in various cell lines, including human amnion cells, lymphoid cells of B cell origin (Fujii et al., 1990), and primary cells from human joint tissue (Huppertz and Chantler, 1991). We established a culture model of persistent MuV infection in a HeLa cell line stably expressing the stress granule protein mCherry-G3BP1 (Ras GTPase-activating protein-binding protein 1) as a marker of cellular stress (Guillen-Boixet et al., 2020) (STAR Methods; Table S1). Our model recapitulated hallmarks of persistent benign infection (Virgin et al., 2009; Walker and Hinze, 1962): cells did not exhibit stress and continued dividing (Figure 1A), while virions released into the medium were capable of infecting naïve cells indicating basal viral replication (Figure 1B). However, when cells were challenged with established conditions of acute or prolonged mild oxidative stress, viral replication increased by 2-4 fold (Figures 1C and 1D). Specifically, transcription levels probed by quantitative PCR of P/V and F genes steadily increased over 6 h under severe stress (1 mM arsenate; As(V)), and that of N stabilized after 3 h, in agreement with a previous report on its self-limiting replication behavior (Doi et al., 2016) (Figure 1C, left). Viral replication, inferred from the genomic RNA level, was similarly up-regulated (Figure 1C, right). Prolonged mild oxidative stress (30 μM arsenite; As(III)) also increased viral transcription and replication, albeit at lower rate or magnitude (Figure 1D). Thus, the increase in the viral activity correlated with the severity of stress. We additionally measured N protein levels in the media as a proxy for viral particle release. While the levels oscillated over 24 h as has been reported for hepatitis C virus (Ruggieri et al., 2012), stress accelerated viral budding by 1.7-fold at 6 h of mild stress compared to the non-stressed cells (Figure S1A). These results substantiate previous reports on the effect of stress on viruses from distant families (Padgett et al., 1998; Piette and Legrand-Poels, 1994), and the closely related MeV (Rima et al., 1977), confirming that cellular stress increases viral transcription, replication, and virion release in our persistent MuV infection model.

**Figure 1.**
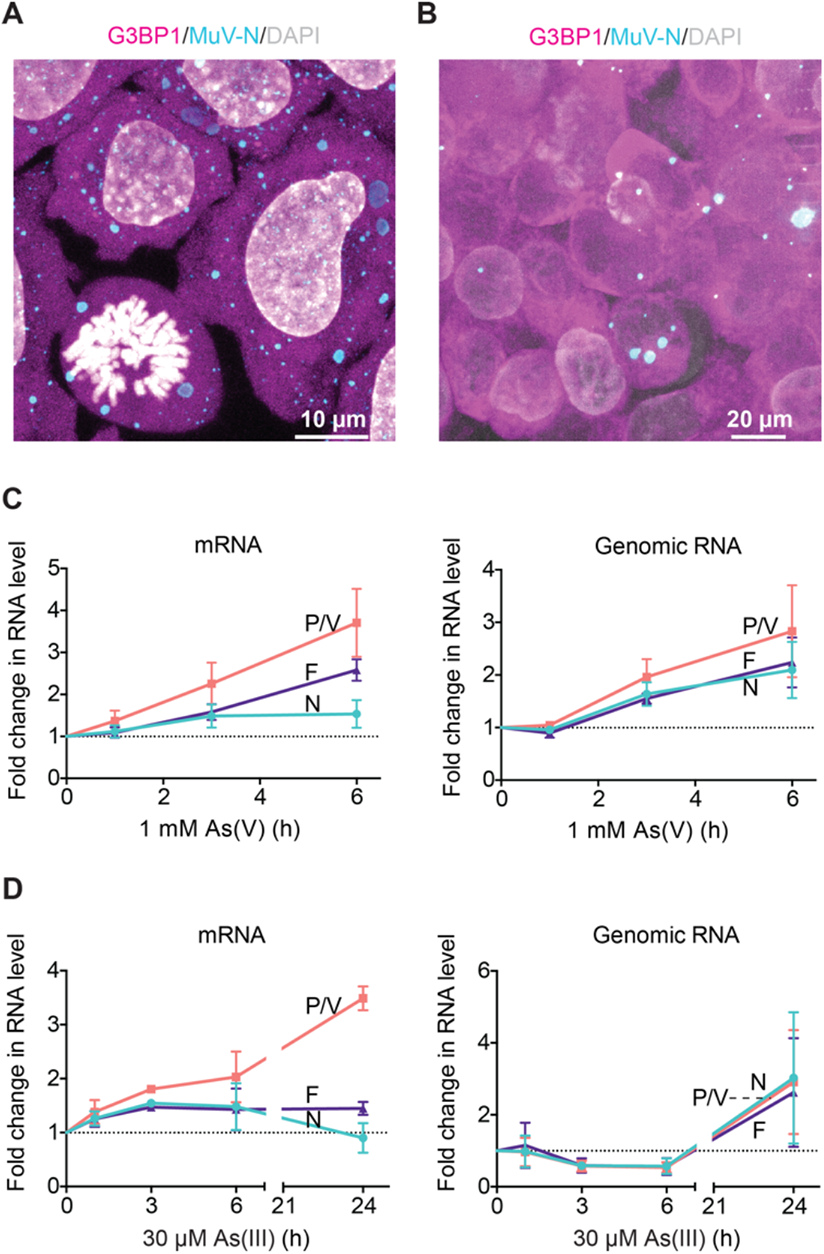
Cell culture model of persistent MuV infection in HeLa and reactivation by stress. (A) Maximum-intensity projections (MIPs) of immunostaining against the MuV-N protein (cyan) in HeLa mCherry-G3BP1 (magenta) cells persistently infected by MuV (HeLa-MuV). DNA was stained with DAPI (grey). Note that cell division does not appear to be affected by persistent infection. (B) MIPs of naïve HeLa mCherry-G3BP1 cells infected with MuV by transfer of medium from a persistently infected culture, imaged at two days post infection. Coloring is the same as in (A). (C and D) MuV transcription (mRNA) and replication (genomic RNA) levels determined by quantitative PCR targeting the N, P/V, and F genes during the time course of 1 mM As(V) (in C: acute) and 30 μM As(III) (in D: mild) stress. Dotted lines: starting levels. Data are mean ± SD (*n* = 3). See also Figure S1.

### MuV replication factories are liquid-like condensates that coarsen under stress

MuV replication occurs in cytoplasmic viral factories. To investigate the cellular mechanisms of stress-induced replication, we monitored changes to the MuV factories using immunofluorescence imaging with antibody against the native viral N protein (Figures 2A, 2B, and S1B-S1D). Both oxidative stress and heat shock elicited stress granules as a part of the cellular response (Guillen-Boixet et al., 2020), and resulted in an increase in viral factory sizes, concurrently with a decrease in their numbers (quantified in Figures 2C-2F). This was reminiscent of the coarsening behavior of liquid-like biomolecular condensates, whereby larger droplets grow at the expense of smaller ones by fusion (Berry et al., 2015) or Ostwald ripening (Zwicker et al., 2015). Indeed, treatment with hexanediol that leads to dissolution of the liquid-like stress granules in our MuV model (Wheeler et al., 2016) also dissolved viral factories to a large extent (40 - 60% in number of large viral factories; Figures 2G and S2A), indicating that MuV factories are liquid-like condensates.

**Figure 2.**
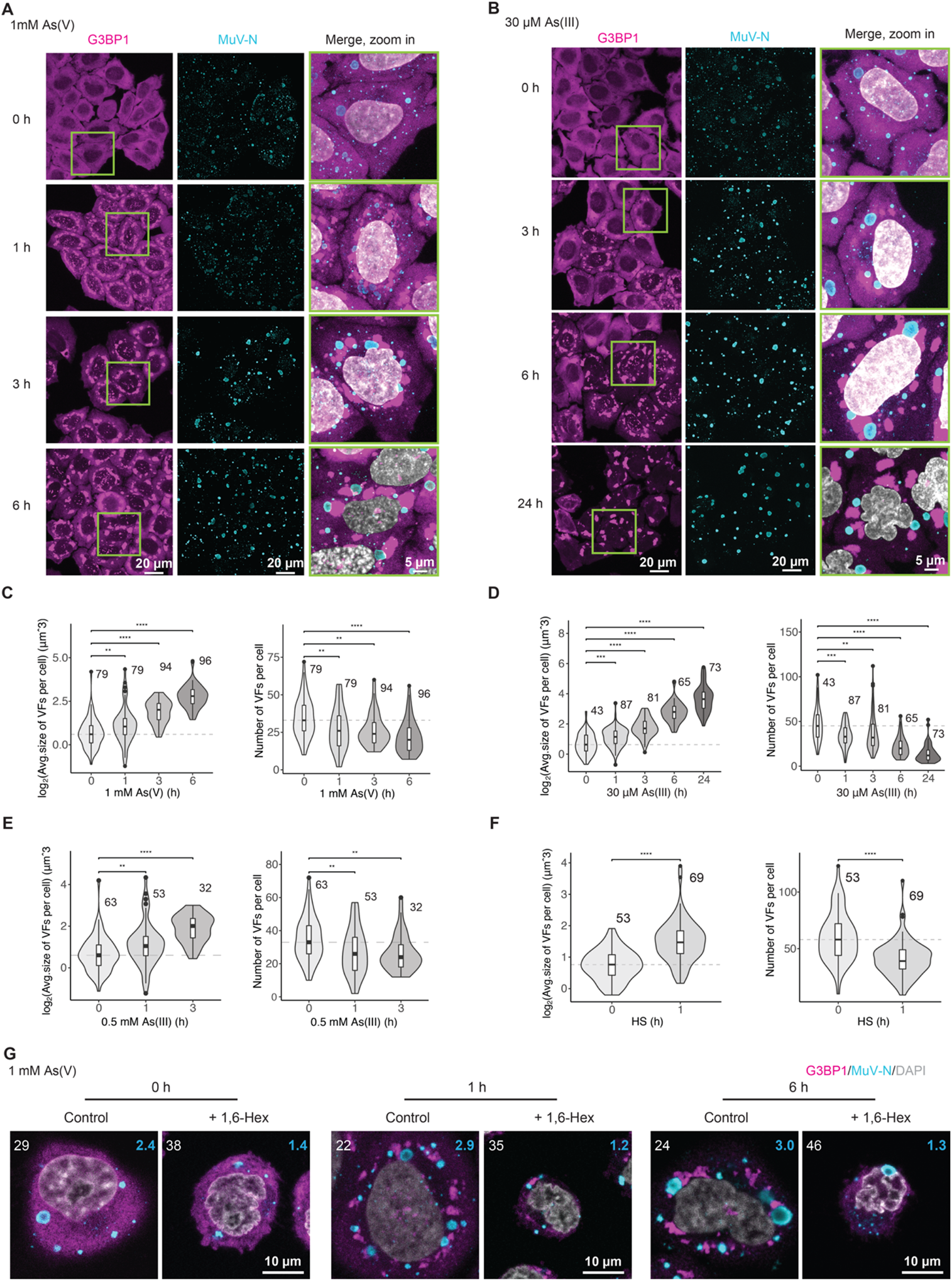
Persistent MuV replication factories are liquid-like condensates that coarsen under stress. (A and B) MIPs of MuV factories (immunostaining against the MuV-N) and stress granules (mCherry-G3BP1) in HeLa-MuV cells during the time course of 1 mM As(V) (in A) and 30 μM As(III) stress (in B). DNA was stained with DAPI (grey). (C and D) Quantification of data in (A and B) for MuV factories (VFs) size and number per cell. Number of cells per condition is indicated. Box center line: median; box bounds: the first and third quartiles; whiskers: values no further than 1.5 times the distances between the first and third quartiles; dashed line: median at the start of stress. Statistical significance is evaluated using Wilcoxon test, \*\**P*<0.01, \*\*\**P*<0.001, and \*\*\*\**P*<0.0001. See also Figure S1B. (E and F) Quantification for VFs size and number per cell over the time course of additional stress conditions: 0.5 mM As(III) and heat shock stress, plotted as in (C). See also Figures S1C and S1D. (G) Effect of 15 minutes 1,6-hexanediol (1,6-Hex) treatment on MuV factories following up to 6 h 1 mM As(V) stress. HeLa-MuV cells were stressed to the indicated time points and then incubated with 1,6-Hex or water as control. Central plane images are shown. The number of cells per condition is indicated on the left corner; number on the right corner is the average number of MuV factories larger than 3 μm in diameter per cell at each condition. See also Figure S2.

Biomolecular condensation is often driven by multivalent interactions, commonly mediated by IDRs (Wang et al., 2018; Zhou et al., 2019). In fact, the highly-transcribed MuV N and P genes encode sequences indicative of disordered regions (Figures S2B and 3A) (Cox et al., 2013; Cox et al., 2009; Pickar et al., 2015). To understand the basis for viral factory condensation, we transfected N and P genes into naïve cells. While N-only transfection resulted in small puncta throughout the cytoplasm (Figure 3B, upper left), P transfection resulted in the formation of spherical cytoplasmic condensates (Figure 3B, bottom left) and its co-transfection with N led to co-condensation of both proteins (Figures 3B, right). Thus, the disordered P protein can drive the condensation of viral factories that change their behavior and coarsen under stress.

**Figure 3.**
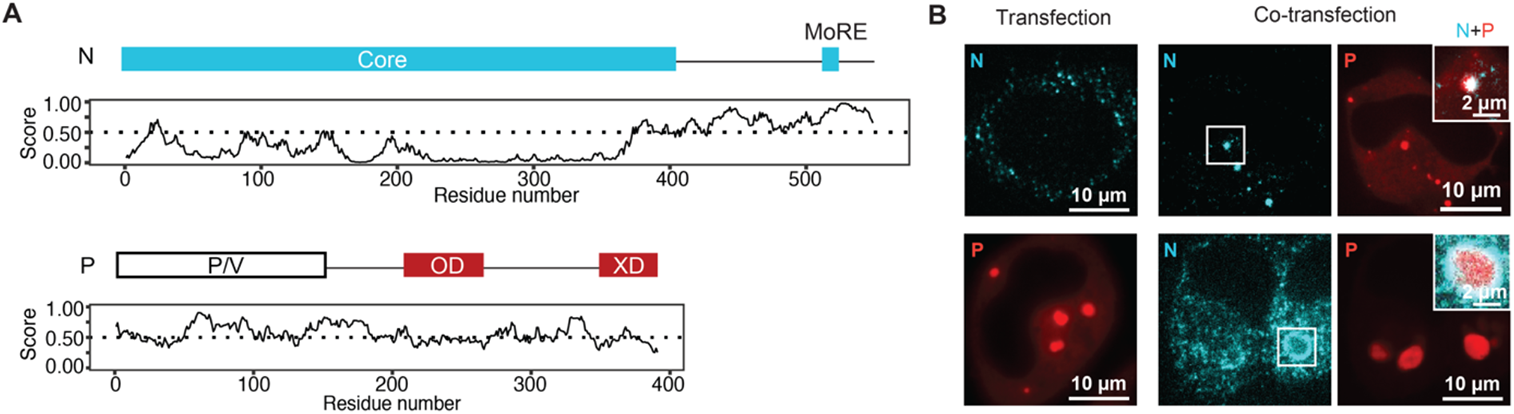
The highly expressed MuV P protein drives condensation. (A) Domain architecture of the viral N and P proteins. Intrinsically disordered regions (IDRs) were predicted with IUPred2A. Residues with scores higher than 0.5 (above the dotted line) are predicted to be disordered. N-Core: core domain of N involved in nucleocapsid assembly; N-MoRE: a short helix that interacts with XD domain of P (a helix bundle); P/V region: the shared region between P and V proteins; P-OD: oligomerization domain of P. (B) Transfection of MuV N (detected by immunostaining) or EGFP-P, or co-transfection of MuV N and P into naïve HeLa mCherry-G3BP1 cells. The size difference of co-condensates of N and P is likely related to the expression levels of the proteins following transient transfection. See also Figure S2.

### Stress alters protein interaction networks within viral factories and induces stabilization of the MuV replication machinery

To elucidate the molecular changes upon stress that lead to induction of the MuV replication, we performed whole-cell quantitative MS (Sridharan et al., 2019). The analysis showed that non-stressed cells expressed high levels of MuV proteins (Figure S3A). Unexpectedly, MuV proteins levels showed modest changes under all examined stress conditions (Figures 4A and S3B; Table S2). However, despite stable total level of P, levels of P phosphopeptides spanning residues 292-328 within its disordered region were significantly elevated during stress (2-fold in Figure 4B and up to 6-fold in Figure S3C, depending on the stress; Table S3). Mapping of the phosphorylation sites of P onto the recently reported structure of viral P-L polymerase complex from the closely-related parainfluenza virus 5 (Abdella et al., 2020) showed that they localize at the P-L interaction interface (Figure 4C). Due to complementary charges at its interface with L, we hypothesized that P phosphorylation may stabilize its interaction with L. In the context of biomolecular condensates, we expected that stabilization of the P-L protein-protein interaction should increase partitioning of L into the MuV factories. We probed this by assessing whole-cell protein solubility using a lysate pelleting assay coupled with quantitative MS (Becher et al., 2018; Sridharan et al., 2019). The solubility profiling revealed that significant proportions of the viral proteins N (∼90%), M (∼80%) and P (∼70%) resided in the insoluble fraction in unstressed cells (Figures 4D, S3D, and S3E; Table S4). This is consistent with N and M propensity to polymerize into structured assemblies (Liljeroos et al., 2011), while P is capable of condensation (Figure 3B). All other viral proteins were predominantly soluble in unstressed cells, suggesting their weak and transient interactions with the core factory proteins N and P (Katoh et al., 2015). L, however, exhibited significant decrease in solubility especially under severe stress (Figures 4D, S3D, and S3E). Such change in solubility suggested increased partitioning of L into the viral condensates. Indeed, sucrose gradient fractionation identified that N, P and L formed a complex that exhibited ∼2-fold enrichment for L under stress (Figures 4E, 4F, and S3F-S3I; Table S5). These data point to a molecular mechanism where stress enhances the interaction of L with the MuV nucleocapsids, likely modulated through phosphorylation of P by host factors (Table S2), and thereby could explain the up-regulation of viral transcription and replication.

**Figure 4.**
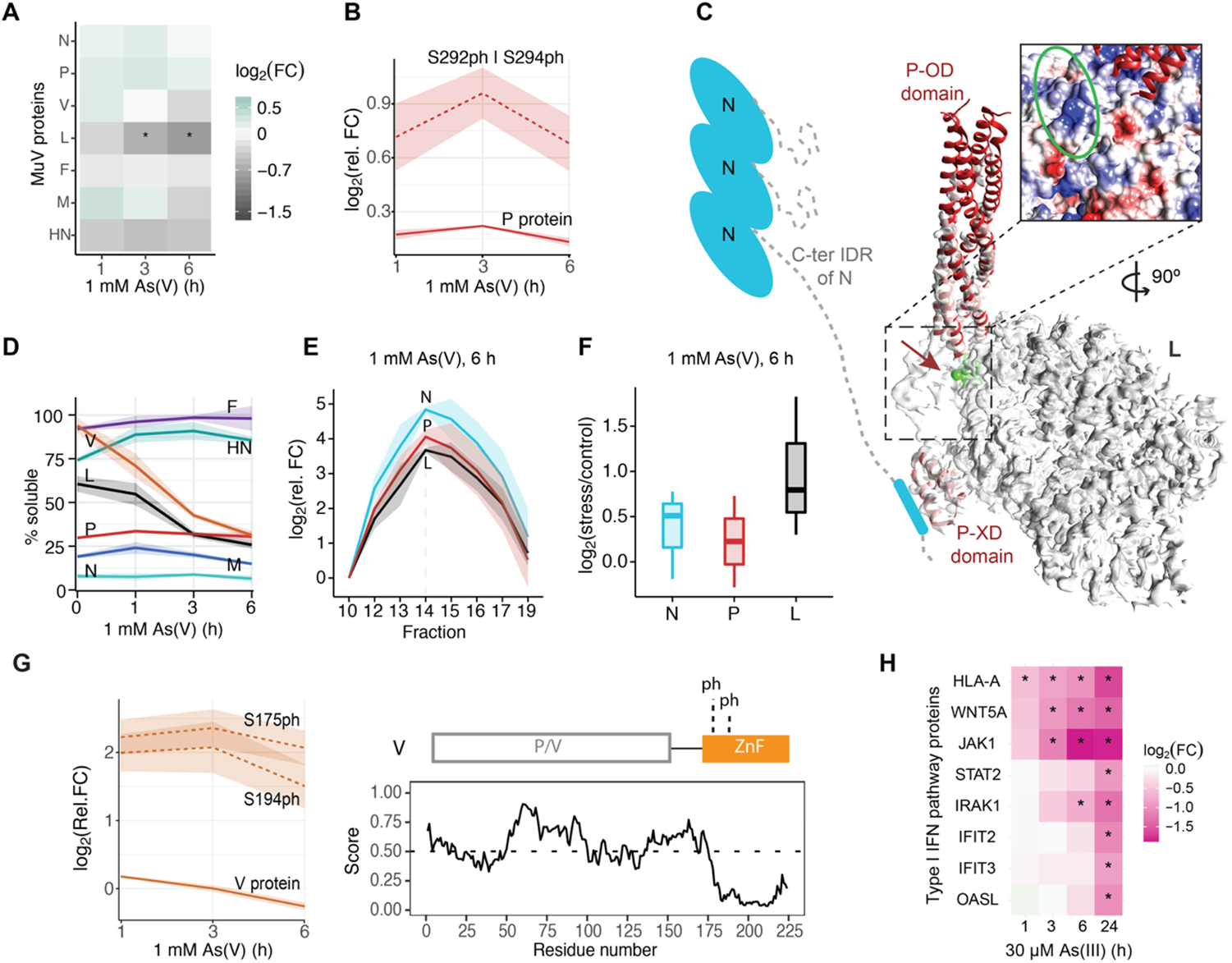
Stress alters the protein networks within viral factories and induces down-regulation of the host immune response. (A) Relative abundance of all MuV proteins in HeLa-MuV cells along the stress time course in comparison to 0 h in log_2_ scale (*n* = 3). * proteins with |log_2_(fold change) | > 0.5 and adjusted p-value < 0.01 (Benjamini-Hochberg method). FC: fold change. (B) Mean relative abundance of P protein (solid line) and its phosphopeptides (dashed line) in HeLa-MuV cells along the stress time course in comparison to 0 h control in log_2_ scale. Shaded areas represent SEM (*n* = 3). (C) P phosphosites (arrow) mapped onto PIV5 P-L complex structure (EMD-21095, PDB 6V85). Inset: view of the predicted interaction interface of P on L (K390 region denoted by green circle), colored according to electrostatic potential (blue to red: positively to negatively charged). Domains are labeled as in Figure 3A. Schematic model of N is shown to indicate the context of P-L complex, with a short helix (MoRE) at the C-terminal IDR of N predicted to interact with the P-XD. (D) Mean solubility profiles (line) and SEM (shaded area) of all MuV proteins (*n* = 2) in HeLa-MuV cells along the stress time course. (E) Mean relative abundance of N, P and L proteins (line) in sucrose gradient fractions with respect to Fraction 10 at stress condition. Apex of co-eluting proteins N, P, and L is observed at Fraction 14. Shaded areas represent SEM (*n* = 3). (F) Distribution of relative abundance of N, P, and L at the apex of stress condition compared to the unstressed control (*n* = 3). (G) Relative fold change of MuV V protein in HeLa-MuV cells along the stress time course relative to 0 h control: unmodified protein (solid line) and phosphopeptides (dashed lines). Lines and shaded areas are mean and SEM (*n* = 3). The domain architecture of V and its IDR prediction are shown on the right. V-ZnF: zinc finger domain of V. Detected phosphorylation sites are indicated. (H) Heat map representation of relative fold change of proteins from type I interferon signaling pathway (HeLa-MuV cells) shows significant down-regulation along the stress time course compared to 0 h control. Asterisks indicate proteins with |log_2_(fold change) | > 0.5 and adjusted p-value < 0.01 (Benjamini-Hochberg method). See also Figure S3.

### Stress increases phosphorylation of the V protein and induces down-regulation of host immune response

In addition to changes that directly affect the viral replication machinery, we found that the co-transcriptional product of P, the V protein, also exhibited significantly increased phosphorylation at sites S175 and S194 (Figures 4G and S3J) and decreased solubility (Figures 4D and S3E). V is known to block the host antiviral response via interaction with STAT1 (Kubota et al., 2002; Yokosawa et al., 2002) or STAT2 (Rosas-Murrieta et al., 2010). Phosphorylation at S194 of V is predicted to localize on its zinc finger domain (Figure 3A), near its interface with STAT1/2 (Angers et al., 2006; Rosas-Murrieta et al., 2010). This suggests a stress-tunable role for V in suppressing host antiviral pathways. In agreement, quantitative MS of the host proteome revealed concomitant changes in the host immune response. While attenuation of the host antiviral defense by degradation of STAT1 has been documented in persistent infection (Kubota et al., 2002; Yokosawa et al., 2002), we additionally found a significant down-regulation of type I IFN signaling pathway proteins, including the mediators Janus kinase 1 (JAK1) and STAT2 from the start of stress (Figure 4H; Table S2). Therefore, post-translational modifications of viral proteins can further support stress-mediated reactivation of viral replication by further suppresion the host defense.

### Stress induces a conformational switch in the MuV nucleocapsids

To structurally assess the consequences of the stress-mediated cellular and molecular changes, we performed cryo-ET on the intracellular viral factories. MuV factories frequently localized in close proximity to stress granules (Figures 2A and 2B). We thus exploited the mCherry-G3BP1 fluorescence to perform 3D targeted cryo-FIB thinning and to locate unlabeled viral factories for cryo-ET (Figure S4A). We observed curved hollow filaments resembling low-resolution maps of MuV nucleocapsids (Cox et al., 2014; Severin et al., 2016) in the viral factories (Figures 5A and 5B) and near the plasma membrane (Figure S4B). 3D tracing of the nucleocapsids allowed quantitative analysis of changes in molecular crowding along the stress course (Figure 5C; Videos S1 to S3). Nucleocapsid volume fractions in the viral factories tripled from the control to 6 h of mild stress (30 µM As(III); Figures 5C, S4C, and 5D). In addition, thin flexible densities extending from the nucleocapsids were abundant under stress conditions (Figures 5A and 5C), potentially marking interaction partners of N. Strikingly, prolonged stress led to a morphological transition from curved to straight nucleocapsids (Figures 5C, 5D and S4D) with the latter reminiscent of aberrant MuV inclusions in myositis patient biopsies (Chou, 1986).

**Figure 5.**
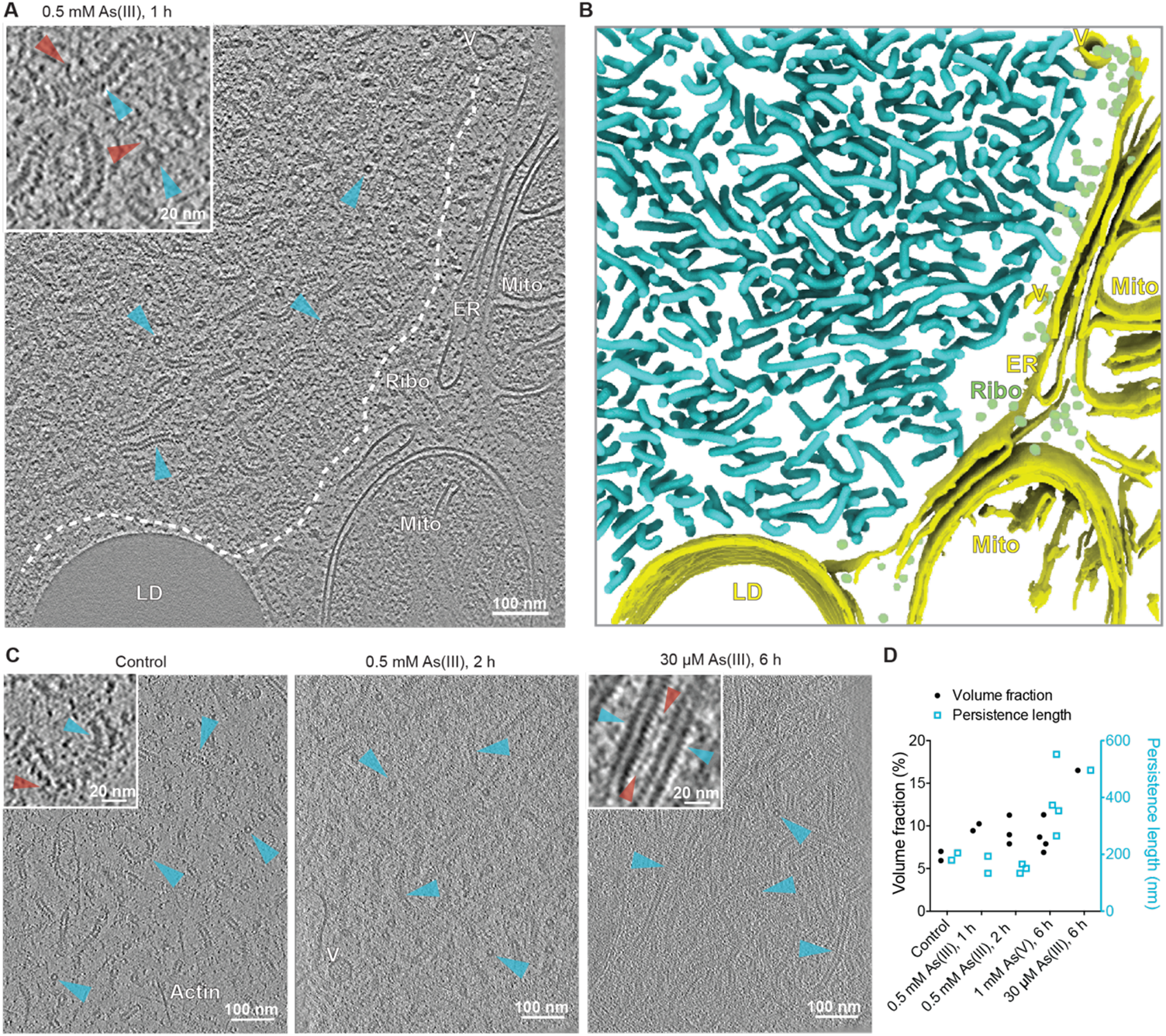
MuV viral factories and nucleocapsids undergo structural changes following stress. (A and B) Tomographic slice (thickness: 6.74 nm) of a MuV factory and the corresponding 3D segmentation. Examples of nucleocapsids are indicated with cyan arrowheads (top and side views, two of each) in (A) and colored the same in (B). Inset: enlarged view. Red arrowheads indicate flexible densities extending from nucleocapsids. LD: lipid droplet; Mito: mitochondrion; ER: endoplasmic reticulum; Ribo: ribosome; V: vesicle. (C) Representative tomographic slices (thickness: 6.74 nm, 6.52 nm, and 6.52 nm – left to right) of nucleocapsids (cyan arrowheads) and associated flexible densities (red arrowheads) at indicated conditions. (D) Volume fractions and persistence lengths of nucleocapsids in tomograms at indicated conditions. See also Figure S4 and Videos S1-S3.

To interpret the morphological transitions in the nucleocapsids under stress, we first generated structures of the authentic MuV nucleocapsids. Obtaining high resolution structures of *Paramyxoviridae* family nucleocapsids has long been impeded by their curved and flexible nature. Available structures were enabled either by digestion or truncation of the flexible C-terminal region of the N protein to induce rigidity (Alayyoubi et al., 2015; Desfosses et al., 2019; Gutsche et al., 2015), or by recombinant N protein expression that resulted in ring-shape or different helical assemblies (Shan et al., 2021). Here, we isolated the authentic MuV nucleocapsids from stressed cells after 1 h of 0.5 mM As(III) (Figures S5A and S5B). The isolated nucleocapsids were curved, similar to their intracellular morphology at early timepoints of stress, but straightened upon addition of the RNA-mimicking molecule heparin (Figure S5A). This change in morphology was similar to the change in nucleocapsids seen inside cells at prolonged stress conditions, suggesting that RNA engagement with the nucleocapsid may contribute to the change from bent to straight conformation.

We obtained two maps from subtomogram averaging on the heparin-straightened nucleocapsids (Figures 6A and S5C-S5E). The two maps exhibited distinctly different helical parameters: a 5.58 nm long-pitch structure constituting the majority class (77.7%) was resolved to 4.5 Å, and a 4.70 nm short-pitch class (21.3%) was resolved to 6.3 Å (Figures 6A, S5E, S6A, and S6B; Table S6). Excluding the most flexible C-terminal IDR of N (144 amino acids) that was not resolved in either map, we built an atomic model for the N protein into the high-resolution majority class based on a homologous structure from parainfluenza virus 5 nucleoprotein (PDB: 4XJN; Figures 6B, S6C, and S6D; Table S6; Video S4). The model showed that the viral RNA genome is accommodated at the nucleocapsid outer surface, within a groove connecting the N-terminal domain (NTD) with the C-terminal domain (CTD) (Figures 6B, 6C, and S6D). The RNA packaging of MuV follows the “rule of six”, *i.e.* six nucleotides per N subunit (Gutsche et al., 2015), and is accommodated by conserved residues from a loop containing Tyr350, Thr351, Arg354 in the upper CTD, and loop and helices containing Lys198, Arg194, Tyr260, Lys180 from the lower NTD (Figure 6F). Interactions between subunits in consecutive turns of the nucleocapsid are mediated by the CTD-arm helix (a region succeeding the CTD) which inserts between the NTDs of two subunits in the upper turn (Figure 6D). Within the same turn, domain swapping between the NTD-arms (a region preceding the NTD) and CTD-arms of consecutive subunits formed extensive contacts at the nucleocapsid lumen (Figure 6D). Compared to the map of the majority class, density of this critical CTD-arm was missing in the map of the minority class indicating its higher flexibility (Figure 6E), as only full-length protein was detected by western blot analysis (Figure S5B). Interestingly, both structural classes existed within the same nucleocapsid (Figure S6E), in line with previous studies on MeV and MuV nucleocapsids (Ke et al., 2018; Shan et al., 2021), which suggested a regulatory role of the C-terminus of N in transcription and replication of *Paramyxoviridae* (Cox et al., 2017; Thakkar et al., 2018). Our two maps thus reflect the structural plasticity of N (Bhella et al., 2004; Cox et al., 2014; Severin et al., 2016; Shan et al., 2021) suggested to contribute to different functional outcomes: genome packaging to protect the viral genome from host antiviral factors versus genome accessibility to promote viral replication. Indeed, tighter packing of subunits in the minority class resulted in an 8.4 Å shift of a loop in the upper subunits towards the lower subunits, resulting in less surface-exposed RNA binding pocket (Figure 6F). The tighter packing also restricted the space available for the flexible C-terminal IDR of N to extend to the nucleocapsid surface (Figures 6F and S6E). In contrast, the loosely-packed structure of the majority class exhibits higher exposure of the RNA on the nucleocapsid surface and higher likelihood for the C-terminal IDR of N protein to interact with partner proteins, *i.e.* P and L, at the nucleocapsid surface. Both factors would contribute to a conformation that can support viral transcription and replication.

**Figure 6.**
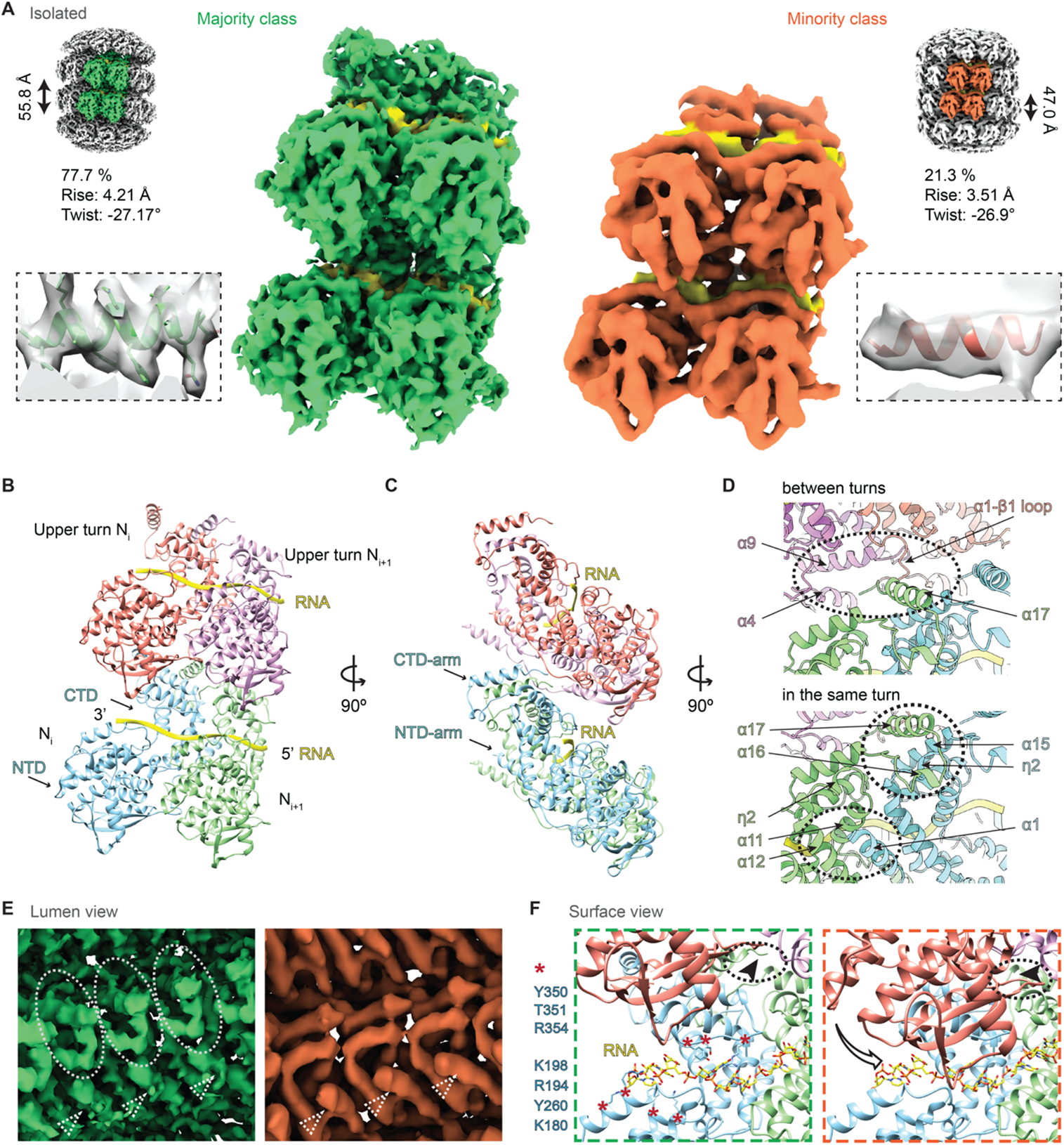
Structures of isolated MuV nucleocapsids reveal two packing modes with variable surface accessibility of RNA and C-terminal IDRs. (A) Two classes of subtomogram averages of extracted nucleocapsids resolved to 4.5 Å (green, left) and 6.3 Å (orange, right). RNA densities are colored in yellow. Four subunits are segmented to illustrate the packing. Zoomed-in views illustrate the quality of the map and model fit. (B and C) Cartoon representation of the atomic model of MuV-N protein (amino acids 3-405) with RNA (yellow) for the majority class. Four subunits are shown with domains labeled. (D) Subunits packing between neighboring turns (upper panel) and within the same turn (lower panel). Circles indicate the packing interfaces. Subunits are colored as in (B). (E) Lumen view comparison of the two classes in (A). CTD-arms of three neighboring subunits (circled) are not resolved in the minority class (orange). Dotted triangles indicate the same NTD-arms in two maps. (F) Surface view comparison of the two classes. Subunits are colored as in (B). RNA is in stick representation and colored by heteroatom. Key residues potentially responsible for RNA binding are labeled (red asterisks on the left). Arrow indicates a shift of the upper subunit towards the lower turn RNA binding pocket in the minority class (orange frame). Dotted circle indicates unoccupied space in the majority class (left) becomes occupied in the minority class (right), which blocks access of the C-terminus (arrowheads) to the nucleocapsid surface. See also Figures S5 and S6.

### Nucleocapsids at prolonged stress adopts a replication accessible state

While a recent study on MuV nucleocapsids also reported that two distinct helical assemblies of the nucleocapsids exist in recombinantly expressed N protein (Shan et al., 2021), the limited biological context of such structures hindered understanding of their relevance to functional states with respect to viral replication. With the structural models for the authentic MuV nucleocapsid at hand, we next examined the conformation of nucleocapsids in the cellular context. Their high curvature at early time points of stress (1 h) provided maps at ∼30 Å only, showing class averages of mixed helical rises (Figures S7A-S7C), in line with non-uniform conformations observed in the isolated nucleocapsids at the same stress timepoint. At prolonged stress, we resolved the straight nucleocapsids to 6.5 Å (30 µM As(III) and 1 mM As(V) for 6 h; Figures 7A and S7D-S7F). A single class was obtained that resembled the loosely-packed majority structure of the isolated nucleocapsids, with clear luminal density for the CTD-arm (Figure 7B) and surface exposed RNA (Figures 7A and 7C). However, additional densities at the nucleocapsid surface were exclusively found in the in-cell map (Figures 7D and 7E). This region was previously speculated to form the interaction interface between N and P, mediated by their largely disordered C-termini (Cox et al., 2014; Jensen et al., 2011). We suggest that the additional density in the in-cell map could be part of the C-terminal IDR of N that becomes rigid due to its interaction with P under prolonged stress, consistent with our observation of abundant flexible densities near the nucleocapsids in the cellular tomograms (Figure 5C, right). Taken together, we conclude that in cells under prolonged stress, the nucleocapsids adopt a homogenous conformation with a surface exposed viral genome that is likely to be more accessible for efficient transcription and replication by L.

**Figure 7.**
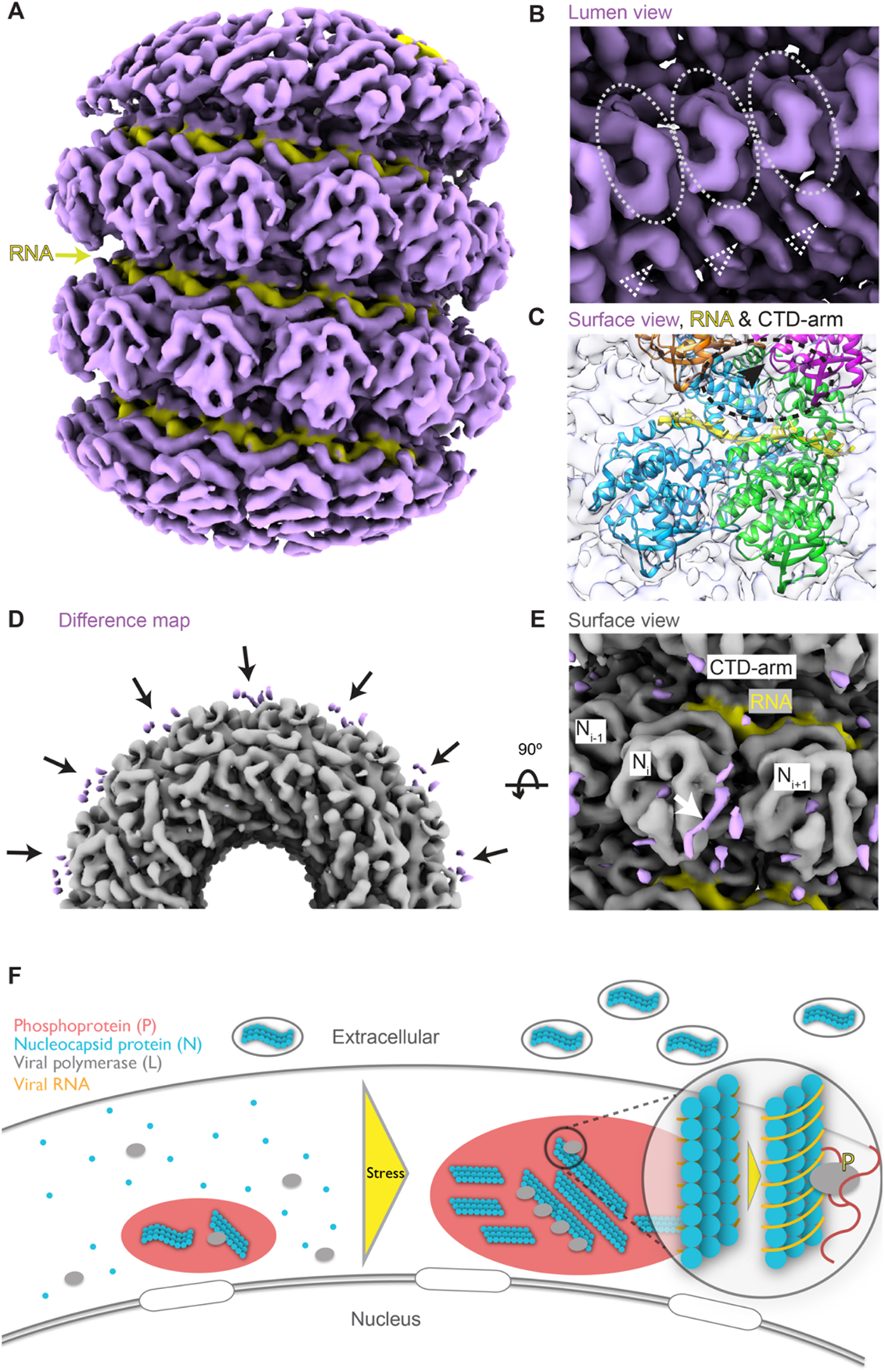
In-cell structure of MuV nucleocapsids at prolonged stress shows an RNA accessible state. (A) Subtomogram average of the in-cell MuV nucleocapsids resolved to 6.5 Å. RNA densities are colored in yellow. See also Figure S7. (B) Lumen view of the in-cell MuV nucleocapsid. CTD-arms of three neighboring subunits are encircled and the NTD-arms are indicated with dotted triangles, similar to the majority class in Figure 6E. (C) Zoom-in surface view of the in-cell MuV nucleocapsid shows accessible RNA (yellow ribbon) and CTD-arm (indicated with black arrowhead and the surrounding region with dotted circle). Subunits are docked and colored as in Figure 6F. (D and E) Cross-section and surface views of the difference map (purple) between the majority class in Figure 6A (grey) and the in-cell nucleocapsids. Black arrows in (D) and white arrow in (E) indicate potential location of the C-terminal IDR at the interface between neighboring subunits. RNA regions are colored in yellow. (F) Proposed model for stress-induced activation of viral replication in phase-separated MuV factories.

## Discussion

Our work demonstrates how a combination of whole-cell proteomics and cryo-ET provides a powerful tool to probe virus replication in the context of biomolecular condensates inside cells and allows us to propose a mechanistic model for stress-mediated reactivation in persistent infection (Figure 7F). During MuV persistent infection, liquid-like viral factories mediate slow replication and release of the virus. Stress triggers posttranslational modification of P by host factors that may promote partitioning of the polymerase machinery into the viral factories. A structural transition in the nucleocapsid, potentially induced by the change in the protein interaction network in the viral factories, allows the RNA to become more accessible to the polymerase. These events, accompanied by a concerted downregulation of the host antiviral response that can be partially modulated by posttranslational modification to the viral V protein, collectively provide an environment that supports up-regulation of viral replication.

An interplay between viral replication and cellular stress has been documented for many viruses in the acute phase of infection following cell entry. Infection by MuV and rabies virus leads to the formation of stress granule via activation of protein kinase R (PKR) stress pathway by the viral RNAs (Hashimoto et al., 2016; Nikolic et al., 2016). In contrast, MeV can interfere with stress granule formation (Okonski and Samuel, 2013) or others, like Zika virus, take advantage of key stress granule proteins or the stress response pathways (e.g. G3BP1) to promote viral gene expression (Bonenfant et al., 2019). Furthermore, acute phase replication of influenza virus is enhanced in cells exposed to arsenite (Amouzougan et al., 2020). However, little is known about how stress affects viruses, especially RNA viruses, at the chronic infection phase. Exceptions are studies of social stress reported to cause reactivation of latent DNA virus, e.g. herpes simplex virus type 1 (Padgett et al., 1998), and of oxidative stress leading to reactivation of the human immunodeficiency virus type 1 (Piette and Legrand-Poels, 1994). We identified a transition from a persistent infection with slow viral replication in the negative-stranded RNA virus MuV into an activated state when cells were challenged with exogenous stress, reflected by elevation in viral transcription, replication, and acceleration of virion budding. While we have observed a spatial association between the MuV replication factories and stress granules, whether a functional exchange between the two compartments remains to be elucidated (Bonenfant et al., 2019; Hashimoto et al., 2016; Nikolic et al., 2016).

Our proposed stress-reactivation model for MuV chronic infection may represent generic molecular mechanisms for reactivation of chronic RNA viruses. First, phosphorylation of the viral P protein likely induces a cascade of changes in the viral replication machinery that accelerates transcription and replication. Second, we observed that the magnitude of increase in replication activity depends on the severity of stress: genomic RNA level under mild As(III) stress was slightly suppressed until 6 h, before showing dramatic up-regulation at 24 h, a phenomenon reminiscent of the initial stage in acute phase infection where transcription is prioritized to generate more material (e.g. polymerase and nucleocapsid protein) for replication (Katoh et al., 2017). Third, immunomodulation by the viral V protein possibly triggered through its phosphorylation provide an additional layer of cross-talk between the stress response and cytokine signaling, exemplified by downregulation of IFN signaling factors, such as JAK1 (Yokota et al., 2003), and further permits viral replication. We observed that viral budding oscillated in both the unstressed and stressed cells over 24 h. In line with this, feedback mechanisms must exist to maintain a long-term balance between the MuV and the host. Finally, the increase in virion release may result in new infection or slow progression of severe chronic diseases (Chou, 1986; Morfopoulou et al., 2017).

By integrating quantitative light microscopy, hexanediol solubilization assays and transient expression of recombinant viral proteins, we identified MuV factories as liquid-like condensates that coarsen under stress. Cryo-ET further showed that MuV factory coarsening is accompanied by an increase in their internal molecular crowding. Owing to enrichment of the nucleocapsids that harbor the viral genome and of the replication machinery under stress, condensation can provide a mechanism to boost viral replication. A similar function was exemplified by the enhanced MeV genome encapsidation within droplets reconstituted *in vitro* with N protein and RNA, as compared to the dilute phase (Guseva et al., 2020). The formation of large viral factories was not only observed in patient biopsies associated with chronic MuV infection, but also during its acute infection (Katoh et al., 2015). Inhibiting the formation of large factories using a dynein inhibitor during MeV infection resulted in significantly decreased viral titer (Zhou et al., 2019). Viral factory coarsening could therefore represent an essential stage in the viral life cycle, regulating viral assembly and budding. On the other hand, the phase-separated MuV replication compartment can shield viral proteins from host antiviral factors such as IFN effectors, or alternatively, sequester IFN mediators into the condensate to block their downstream antiviral effects as demonstrated by the lowered solubility of V protein and JAK1 in our MS data of stress conditions (Table S4). Unlike previous studies on recombinant MeV (Zhou et al., 2019), our study did not identify a liquid to gel-like phase transition associated with the increased size of MuV viral factories, as probed by the hexanediol solubilization assay. Establishment of persistent infection with genetically-tagged authentic viruses will be needed for monitoring such dynamic changes in the condensate material properties and investigating their functional consequences. Interestingly, a recent study highlights the detrimental effect of viral factory solidification in human respiratory syncytial virus on viral replication (Risso-Ballester et al., 2021).

We identified MuV P protein as the potential driving force for condensate formation, and that it can recruit N protein to form condensates resembling viral replication factories. Consistent with these observations are previous findings in MeV showing that the interaction between the C-termini of N and P is critical for their phase separation (Zhou et al., 2019). Although the sequences of the C-termini of N and P in MeV and MuV are distinct, both appear to be disordered. The rabies virus P protein was also found to be critical for phase separation via its dimerization domain and the flanking IDR (Nikolic et al., 2017). In vesicular stomatitis virus, N, P, and L proteins are all required for formation of condensates, while replication activity is not needed (Heinrich et al., 2018). Interestingly, region-specific genomic RNA (Iserman et al., 2020) and the membrane protein (Lu et al., 2021) of SARS-CoV-2 can independently induce phase separation of its N protein, forming annular structures in which the membrane protein coats the outside of an N + RNA condensate. This further leads us to speculate that in our MuV model, increased viral RNA levels following both upregulated transcription and replication under stress may also modulate the phase behavior of the viral condensates. Another layer for regulation of the properties and dynamics of viral condensates is achieved through phosphorylation. This was reported for MeV (Zhou et al., 2019) and SARS-CoV-2 (Carlson et al., 2020). It is plausible that the phosphorylation identified in our study on the IDRs of both P and V proteins may act as such a fine-tuning factor. The levels of many kinases and phosphatases changed under stress (Table S2). However, the enzymes responsible for the phosphorylation of P and V need further investigation, with the activity changes of enzymes also taken into account. Altogether, these studies reveal that RNA viruses evolved multiple mechanisms to ubiquitously form condensate-mediated viral replication factories.

On the structural level, MuV nucleocapsids are templates for both viral transcription and replication. Their conformational flexibility has been reported for other contagious viruses with negative-stranded RNA genome such as MeV (Bhella et al., 2004; Schoehn et al., 2004) and respiratory syncytial virus (Tawar et al., 2009), which was hypothesized to contribute to regulation of the balance between transcription and replication (Bhella et al., 2004). A previous study on MeV shows clearly that two distinct helical assemblies of the nucleocapsids exist in released virions (Ke et al., 2018), but the functional implication was not addressed due to limited resolution of the structures. In this study, we identified two conformations that coexist in the authentic MuV nucleocapsids isolated from cells after 1 h of cellular stress, which differ in the flexibility of the CTD-arm region of N protein and surface exposure of the viral genome. The ratio of the two conformations may therefore be important for fine-tuning viral activity at persistent infection and for the reactivation under stress. Interestingly, our map of nucleocapsids *in situ* from prolonged stress conditions resolved to high detail represents a genome-accessible conformation, in line with increased viral transcription and replication detected in our PCR quantification. Additional densities on the nucleocapsid surface revealed in this in-cell map in comparison to the extracted nucleocapsids are suggested to be the C-terminus of N or its partner protein P, consistent with previous computational simulations (Jensen et al., 2011) or low-resolution models (Cox et al., 2014), further supporting active state of nucleocapsids at the stress conditions. Finally, the straight morphology of MuV nucleocapsids observed in our data mimics their appearance in chronic myositis, providing a link of such conformation of nucleocapsids to pathology.

In summary, our integrated structural cell biology study illustrates that stress-induced subtle changes in the phosphorylation state of viral proteins and fine structural changes in the nucleocapsid seem sufficient to disturb the delicate balance between the virus and host cell during persistent infection. It is tempting to speculate that such molecular switches induced by stress may have broad implications across a diverse family of viruses that establish chronic infection in the human host, and for the diseases they elicit.

## Supporting information

Key resources table

## Acknowledgements

We thank EMBL core facilities, including the EMCF, ALMF, Genecore, central IT and especially Thomas Hoffman for computational support, Wim Hagen, Felix Weis, and the EMBL cryo-EM platform. The cryo-confocal microscope was developed in collaboration with Leica Microsystems, and we thank Jan De Bock and Martin Schorb for invaluable input and support. We thank the Mahamid group members for fruitful discussions, especially Mauricio Toro-Nahuelpan for help in grid preparation, Liang Xue and Edoardo D’Imprima for help in data analysis. We are grateful to Giulia Zanetti, Daniel Castano-Diez, and Beata Turonova for advice on subtomogram averaging strategies, to Dimitry Tegunov for developing helical symmetry functionality in M, and to Jan Kosinski, Martin Beck and Petr Chlanda for critical reading of the manuscript. H.K.H.F was supported by a fellowship from the EMBL Interdisciplinary Postdoctoral Program (EI3POD) under Marie Skłodowska-Curie Actions COFUND (664726). J.M. acknowledges funding from the EMBL and the European Research Council starting grant (3DCellPhase^−^ 760067).

## Author contributions

X.Z. and J.M. conceived the project. X.Z. collected and analyzed LM, cryo-confocal, negative-staining EM data, prepared samples for PCR, western blot, MS, transcriptome sequencing and cryo-ET, collected and processed cryo-ET data and built the model, with supervision from J.M.; S.S. conducted MS experiments and analyzed the data with supervision from M.M.S.; I.Z. maintained the cell line, preformed western blot, prepared plasmids, and assisted LM experiments; C.E.O. conducted quantitative PCR experiments and analyzed the data; C.C. and T.P. assisted in the cryo-ET data analysis of the intracellular and isolated nucleocapsids; H.K.H.F., under the supervision of C.W.M., assisted with sucrose gradient fractionation and scripts for CLEM; I.P. provided the cell line and helped with LM; A.A.H. advised on the project, provided additional interpretations and implications; X.Z., J.M. and S.S. made figures; X.Z., S.S, A.A.H, M.M.S. and J.M. wrote the manuscript with input from all authors.

## Declaration of interests

A.A.H. is a founder and scientific advisory board member of Dewpoint Therapeutics and a founder of Caraway Therapeutics. I.P. is an employee of Dewpoint Therapeutics. All authors declare no competing interests.

## STAR METHODS

### KEY RESOURCES TABLE

#### RESOURCE AVAILABILITY

##### Lead contact

Further information and requests for resources and reagents should be directed to and will be fulfilled by the lead contact, Julia Mahamid.

##### Materials availability

No new unique reagents were generated in this work.

##### Data and code availability

Structures for initial model building and analysis of interaction interfaces were obtained from the Protein Data Bank (PDB) with accession codes 4XJN (PIV5 nucleocapsid-RNA complex) and 6V85 (PIV5 L-P complex), and the Electron Microscopy Data Bank (EMDB) under accession code EMD-21095 (EM map of PIV5 L-P complex). Tomograms generated in this study are deposited in EMDB under accession codes EMD-13165 (viral factory at non-stressed condition), EMD-13166 (viral factory at 1 h of 0.5mM As(III) stress), and EMD-13167 (viral factory at 6 h of 30 µM As(III) stress). Density maps are deposited under accession codes EMD-13133 (majority class, isolated), EMD-13136 (minority class, isolated), EMD-13137 (straight cellular nucleocapsids). The associated model is deposited in the PDB under accession code 7OZR, and raw micrographs in the Electron Microscopy Public Image Archive under accession code EMPIAR-10751 (isolated nucleocapsids). The cryo-EM maps and atomic model deposited to the public repositories will be released upon publication. Mass spectrometry data are deposited in PRIDE. Data are available via ProteomeXchange with identifier PXD026799.

#### EXPERIMENTAL MODEL

The HeLa cell line stably expressing mCherry-G3BP1 by bacterial artificial chromosome tagging (Poser et al., 2008) was previously described (Guillen-Boixet et al., 2020). Cells were cultured in Dulbecco’s modified Eagle’s medium (DMEM, high glucose, GlutaMAX Supplement, Gibco) supplemented with 10% fetal bovine serum, 50 U/ml Penicilin-Streptomycin and 2 µg/ml Blasticidin S (Gibco), and maintained using standard procedures. We accidently encountered the persistent MuV infection (strain Enders, single nucleotide variations in Table S1) in our cell culture (HeLa-MuV cells). Establishment of persistently infected cell line was performed as reported previously (Fujii et al., 1990).

#### METHOD DETAILS

##### Stress treatment and confocal light microscopy

For fluorescent microscopy, HeLa-MuV cells (mCherry-G3BP1) were seeded on 35-mm dishes with glass bottom the day before stress treatment. Cells were stressed by replacing the culture medium with that containing either 0.5 mM sodium arsenite (referred to as As(III)) for up to 3 h, 1 mM potassium arsenate (referred to as As(V)) for up to 6 h, 30 µM As(III) for up to 24 h, or transferring the cell culture dishes to an incubator set to 43 °C for 1 h of heat shock. Samples of all time points in particular stress condition were fixed at once with 4% paraformaldehyde (PFA) pH 7.4 in phosphate-buffered saline (PBS; 2.67 mM KCl, 1.5 mM KH_2_PO_4_, 137 mM NaCl, and 8.1 mM NaH_2_PO_4_, pH 7.4) for 10 min, after rinsing twice with PBS. Samples were rinsed with cold PBS and permeabilized with 0.2% Trion X-100 for another 10 min. Samples were blocked with 1% bovine serum albumin (BSA) for 30 min, incubated with mouse monoclonal antibody against MuV N protein (1:200; clone 8H4; Abcam, ab9876) diluted in 1% BSA for 1 h at room temperature and then with goat anti-mouse IgG secondary antibody Alexa Fluor 488 (1:2,000; highly cross-adsorbed; Invitrogen, A-11029). The samples were finally immersed in ProLong Gold Antifade Mountant formulated with the DNA stain DAPI (Thermo Fisher Scientific) and sealed. Confocal imaging was performed on a Zeiss LSM780 inverted microscope with a 63× 1.40 NA oil immersion objective and a GaAsP spectral detector. Settings were kept the same for imaging samples of all time points in one experiment. Z-stack images were segmented using the Imaris software (Oxford Instruments) using the Surface module. Surfaces of viral factories were generated by absolute intensity thresholding with a fixed value for the anti-N staining signal in each dataset (values optimized for different datasets) and surfaces of nuclei by background subtraction auto-thresholding of the DAPI staining signal, separately. Distances between viral factories and nuclei were calculated for each image stack with MATLAB (MathWorks) scripts to assign viral factories to individual cells. Volumes and numbers of viral factories per cell were then used for generating density plots with RStudio scripts.

All light microscopy figures were prepared with FIJI (Schindelin et al., 2012). Maximum intensity projected images are shown for Z-stack images.

##### Infection assay

To test if the MuV virions released during persistent infection are infectious, naïve HeLa mCherry-G3BP1 cells (not infected by MuV) were used for the infection assay. One day before the assay, naïve HeLa mCherry-G3BP1 cells were seeded on one 8-well slide (Ibidi, µ-Slide) to reach 80∼90% confluent on the day of infection. Virions were collected from culture medium of HeLa-MuV cells after removing cell debris by centrifugation at 2200*g*, 4 °C for 15 min. The original virion stock was serially diluted to 10^−1^, 10^−2^, 10^−3^, 10^−4^, 10^−5^, 10^−6^ solutions with DMEM medium (no fetal bovine serum or Penicilin-Streptomycin). After removing the culture medium of the pre-seeded naïve HeLa mCherry-G3BP1 cells and rinsing the cells twice with PBS, 100 µL of the serially diluted solutions and the original virion stock were added separately to each well and incubate for 2 h. During incubation, shake the slide every 30 min. After replacing the infection solution with complete culture medium, culture the cells for 48 h and fix them for checking viral factory formation by immunostaining as described above.

##### Hexanediol treatment

Hexanediol treatment was done for assessing the liquid-like properties of viral factories at 1 mM As(V) stress conditions. Culture medium was replaced with medium containing 1 mM As(V) as before for 1 h or 6 h to induce stress. Together with the unstressed cell samples as 0 h control, cells were either immediately fixed as control for each time point, or incubated with 3.5% 1,6-hexanediol (Sigma-Aldrich) or water for 15 to 30 min and then fixed. Immunostaining and imaging were done as described above. Central slice images were acquired. 2D segmentation was similarly done using the surface module in Imaris software. The number of viral factories larger than 7.07 μm^2^ (diameter over 3 μm assuming a sphere) per cell was counted for each condition.

##### Transfection

Transfection was done with the untagged N and mEGFP (monomeric enhanced green fluorescent protein) tagged P plasmids into naïve HeLa mCherry-G3BP1 cells (not infected by MuV). The genes were synthesized by GeneArt Gene Synthesis (Thermo Fisher Scientific) both in a pcDNA3.1 vector and the mEGFP gene was then inserted at the N-terminus of P gene. Transfection was done on pre-seeded HeLa cells (about 70% confluency) with the jetPRIME Transfection Reagent (Polyplus) following standard protocol. At 48 h after transfection, cells were fixed and immunostained as described above. Imaging was performed on a PerkinElmer spinning disk confocal microscope with a 63× 1.40 NA oil immersion objective and a Hamamatsu EMCCD camera. Central slice images were acquired.

##### Quantitative PCR

0.5 to 1 million cells per condition and time point were used for isolating total RNA. The extraction was performed according to manufacturer’s instructions using the RNeasy Micro Kit (Qiagen). Strand-specific reverse transcription of genomic RNA (gRNA), anti-genomic RNA (cRNA) and mRNA was performed with the 3’-UTR genomic forward primer, L-gene antigenomic reverse primer (Table S7), and oligo d(T)_23_ primer (New England Biolabs) for transcripts. For quantitative comparison between samples, cDNA was also generated for the cellular control gene RNase P using a gene specific reverse primer (Table S7). 1 µg of RNA was used to generate specific cDNAs for each condition, time point and target gene separately. Synthesis was performed according to manufacturer’s instructions using the ProtoScript II Reverse Transcriptase Kit (New England Biolabs).

Real-time quantitative PCR (qPCR) assay was performed using TaqPath qPCR Master Mix, CG (Thermo Fisher Scientific), target-specific primers and Taqman probes (Table S7). 20× primer probe mixes of each target gene were prepared with 150 nM forward primers, 150 nM reverse primers and 100 nM Taqman probes. Genome, and antigenomic cDNAs were mixed with the cellular control gene RNase P cDNA for each condition and time point at a dilution of 1:10. Transcript cDNAs generated with the oligo d(T)_23_ primer were diluted at 1:10 for each condition and time point. The qPCR was set up in 10 µl total mixture comprised of 5 µl Master Mix (2×), 0.5 µl N-gene primer-probe mix, 0.5 µl P-gene primer-probe mix, 0.5 µl F-gene primer-probe mix, 0.5 µl RNaseP control gene primer-probe mix, 2 µl H_2_O, and 1 µl cDNA mixes (1:10 dilution). Within each plate, standard curves were included at the dilution range of 1:10 to 1:10,000 with the non-treated control condition of each biological replica. qPCR was performed using the Roche LightCycler96 cycling conditions: 95 °C for 20 sec for polymerase activation followed by 45 cycles of 95 °C for 15 sec denaturation and 60 °C for 60 sec annealing and extension. The Taqman probes were detected with the filter sets 470/514 nm for FAM, 533/572 nm for HEX, 577/620 nm for LC610, and 645/697 nm for Cy5.

Fold changes were calculated relative to the non-treated 0 h time point of each condition and biological replica. Data was normalized against RNase P internal control. qPCR efficiency values were calculated for each biological replica and included for fold change calculations. The average was calculated for three independent biological replicas to obtain the fold change of genomic and anti-genomic viral load, and transcriptional mRNA levels for each target gene. Data were analyzed with GraphPad Prism 6.0c.

##### Transcriptome sequencing

HeLa-MuV cells after ∼1 h of 0.5 mM As(III) were used for isolating total RNA according to manufacturer’s instructions using the RNeasy Micro Kit (Qiagen). Briefly, RNA integrity was checked using the RNA Nano 6000 Assay Kit of the Bioanalyzer 2100 system (Agilent Technologies), and concentration was measured with Qubit RNA Assay Kit in Qubit 2.0 Fluorometer (Life Technologies). Stranded mRNA-Seq libraries were prepared from 250 ng of total RNA using the NEBNext Ultra II Directional RNA Library Prep Kit for Illumina (New England Biolabs) implemented on the liquid handling robot Beckman i7. The protocol starts with the isolation of mRNA from total RNA by a polyA selection with oligo d(T) beads, followed by fragmentation and priming of mRNA with random primers to generate double-stranded cDNA fragments. Subsequently, adaptors were ligated to cDNA fragments, which were then amplified with 13 PCR cycles and purified with SPRIselect beads (Beckman Coulter). Fragmentation time purification steps were optimized to allow the selection of larger fragment sizes (>200 nt). Size distribution of the final library was assessed on Bioanalyzer with a DNA High Sensitivity kit (Agilent Technologies), and concentration was measured with Qubit DNA High Sensitivity kit in Qubit 2.0 Fluorometer. Final libraries were pooled in equimolar amounts and loaded at 2 pM solution on the Illumina sequencer NextSeq 500 MID output and sequenced uni-directionally, generating ∼150 million reads per run, each 155 bases long.

Sequencing reads were aligned using BWA-MEM (v0.7.17-r1188) with default parameters to the Enders strain of Mumps virus (GU980052) (Young et al., 2009). Duplicated reads were then filtered out using MarkDuplicates function in Picard tool (v2.9.0-1-gf5b9f50-SNAPSHOT). FreeBayes (v1.1.0-3-g961e5f3) was used to call single nucleotide variations compared to the Enders strain sequence. Finally, the variants were filtered based on their qualities (> 20).

##### Western blot

HeLa-MuV cells were seeded ∼24 h prior to the start of stress treatment in T-25 flasks for each sample. Stress treatment was performed by adding 100 uL of 1.5 mM sodium arsenite solution into each T-25 flask (final concentration of 30 µM in 5 mL culture medium each) and gently mixing it with culture medium. Samples were collected at four time points-0 h, 6 h, 12 h, 24 h, for both stressed and control samples. First, the culture medium of each was separately collected for subsequent enrichment of the released N protein. Then, cell pellet samples were collected *via* direct lysis by adding 1% SDS dissolved in PBS, with 1.5 mM MgCl_2_, EDTA-free protease inhibitor cocktail (cOmplete, Roche) and 1 mM PMSF. 0.3 µl of Benzonase (250 U/µl, Millipore) was added to each flask and the flasks were shaken on a shaking platform (Eppendorf Thermomixer Compact) at 750 RPM for 15-20 min until the jelly substance disappeared. The cell lysate was then recovered from the flasks, adjusted to the same volume, heated at 95 °C for 5 min and frozen for further use. Processing of the culture medium containing the released viral proteins was started by centrifugation at 2,200*g*, 4 °C for 15 min to remove cell debris. Then, the supernatant was mixed with 4× concentration solution (40% PEG 6,000, 100 mM HEPES, pH 7.5, and 2 M NaCl, with EDTA-free protease inhibitor cocktail and 1 mM PMSF) thoroughly and left at 4 °C. After ∼24 h of precipitation, the samples were centrifuged at 4,300*g*, 4 °C for 30 min. After careful removal of the supernatant, a white disk was seen at each tube bottom, and recovered with the same lysis buffer as for cell pellets. Volumes of these samples containing the released viral proteins were also adjusted to the same for all samples, heated at 95 °C for 5 min and frozen for further use.

Total protein concentration of each cell lysate sample was determined by Bradford assay with Protein Assay Dye (Bio-Rad) on a UV/Visible spectrophotometer (Ultrospec 2100 pro, Amersham Biosciences). For western blot analysis, volume corresponding to ∼20 µg total protein was taken for each cell lysate sample and supplemented to the same volume for all samples before the loading buffer with reducing agent was added. The samples containing the released viral proteins were also volume adjusted according to the total protein concentration in their respective cell lysate sample so that the comparison will be performed with the same total intracellular protein level among all time points, for both the stressed and control samples. Proteins were separated using NuPAGE 4-12% precast Bis-Tris protein gels in MOPS SDS running buffer (Invitrogen), transferred onto Immobilon-P PVDF membranes (Millipore), and analyzed with mouse monoclonal antibodies against MuV-N (1:1,000; clone 8H4; Abcam, ab9876) and glyceraldehyde-3-phosphate dehydrogenase (GAPDH, 1:4,000; clone 6C5; Santa Cruz, sc-32233) as loading control. Goat anti-mouse IgG (H/L):HRP (1:1,000; multi species absorbed; Bio-Rad, STAR117P) and ECL western blot detection reagents (Cytiva) were then used for detection with a ChemiDoc Imager (Bio-Rad). Intensity quantification was done in Image Lab (v6.1; Bio-Rad). To calculate the intracellular levels of MuV-N over time (fold change), the intensity of N in cell lysate samples was normalized to the intensity of the respective loading control (GAPDH) and shown as relative to 0 h. To analyze the released levels of N over time (fold change), the intensity of MuV-N in samples containing the released viral proteins was divided by the intensity of MuV-N in the respective cell lysate sample and shown as relative to 0 h. Data were analyzed with GraphPad Prism 6.0c.

##### Mass spectrometry (MS) cell lysate preparation

###### Whole cell lysate preparation

Cells were stressed as described above. At the indicated time points, cells were detached by trypsinization, washed and pelleted twice with PBS. For total proteome analysis, 0.5 million cells per condition were lysed with 100 µl of PBS containing 1.5 mM MgCl_2_, 1% SDS and 0.25 U/µl Benzonase and incubated at room temperature for 15 min. The protein concentration in cell lysates were determined using a BCA assay. Cell lysate volumes corresponding to 5 µg total protein from different conditions were utilized for MS sample preparation.

###### Solubility proteome profiling (SPP) lysate preparation

One cell pellet containing 1 million cells was frozen per time point following cellular stress (as described above). The cell pellets were thawed on ice and lysed (3 freeze thaw cycles) after re-suspending each one in 100 µl of PBS containing protease inhibitors (Roche), 0.8 % NP40, 1.5 mM MgCl_2_ and phosphatase inhibitor (Roche). The lysate was split into two 50 µl aliquots. One aliquot was spun at 100,000*g*, 20 min at 4°C and the supernatant was retrieved. This represented the “soluble-NP40” fraction of proteins. The second aliquot was further solubilized with SDS (final concentration 1%). This represented the “total-SDS” proteome. Both aliquots were incubated with Benzonase (final concentration 0.25 U/µl). The protein concentration of total proteome fractions was determined with BCA assay. Volumes of total lysate corresponding to 5 µg protein from different conditions were calculated. Equal volume of the soluble-NP40 samples to its corresponding total lysate was utilized. Both soluble and total lysate of each condition over the time course was multiplexed as a single MS experiment.

##### MS sample preparation

MS sample preparation and measurements were performed as described (Mateus et al., 2020; Sridharan et al., 2019). Protein digestion was performed using a modified SP3 protocol (Hughes et al., 2019; Mateus et al., 2020). 5 µg of proteins (per condition) were diluted to a final volume of 20 µl with 0.5% SDS and mixed with a paramagnetic bead slurry (10 µg beads (Sera-Mag Speed beads, Thermo Fischer Scientific) in 40 µl ethanol). The mixture was incubated at room temperature with shaking for 15 min. The beads now bound to the proteins were washed 4 times with 70% ethanol. Proteins on beads were reduced, alkylated and digested using 0.2 µg trypsin, 0.2µg LysC, 1.7 mM TCEP and 5 mM chloroacetamide in 100 mM HEPES, pH 8. Following an overnight incubation, the peptides were eluted from the beads, dried under vacuum, reconstituted in 10 µl of water and labelled with TMT-10 or TMT-16plex reagents and dissolved in acetonitrile at 1:15 (peptide:TMT weight ratio) for one hour at room temperature. The labelling reaction was quenched with 4 µl of 5% hydroxylamine and the conditions belonging to a single MS experiment were pooled together. The pooled sample was desalted with solid-phase extraction after acidification with 0.1 % formic acid. The samples were loaded on a Waters OASIS HLB µelution plate (30 µm), washed twice with 0.05% formic acid and finally eluted in 100 µl of 80% acetonitrile containing 0.05% formic acid. The desalted peptides were dried under vacuum and reconstituted in 20 mM ammonium formate. The samples were fractionated using C18-based reversed-phase chromatography running at high pH. Mobile phases constituted of 20 mM Ammonium formate pH 10 (buffer A) and acetonitrile (buffer B). This system was run at 0.1 ml/min on the following gradient: 0% B for 0 – 2 min, linear increase 0 - 35% B in 2 – 60 min, 35 – 85% B in 60 – 62 min, maintain at 85% B until 68 min, linear decrease to 0% in 68 – 70 min and finally equilibrated the system at 0% B until 85 min. Fractions (0.2 ml) were collected between 2 – 70 min and every 12^th^ fraction was pooled together and vacuum dried.

##### LC-MS-MS measurement

Samples were re-suspended in 0.05% formic acid and analyzed on Q Exactive Plus or Orbitrap Fusion Lumos mass spectrometers (Thermo Fischer Scientific) connected to UltiMate 3000 RSLC nano system (Thermo Fisher Scientific) equipped with a trapping cartridge (Precolumn; C18 PepMap 100, 5 μm, 300 μm i.d. × 5 mm, 100 Å) and an analytical column (Waters nanoEase HSS C18 T3, 75 μm × 25 cm, 1.8 μm, 100 Å) for chromatographic separation. Mobile phase constituted of 0.1% formic acid in LC-MS grade water (Buffer A) and 0.1% formic acid in LC-MS grade acetonitrile (Buffer B). The peptides were loaded on the trap column (30 μl/min of 0.05% trifluoroacetic acid in LC-MS grade water for 3 min) and eluted using a gradient from 2% to 30% Buffer B over 2 h at 0.3 μl/min (followed by an increase to 40% B, and a final wash to 80% B for 2 min before re-equilibration to initial conditions). The outlet of the LC-system was directly fed for MS analysis using a Nanospray-Flex ion source and a Pico-Tip Emitter 360 μm OD × 20 μm ID; 10 μm tip (New Objective). The mass spectrometer was operated in positive ion mode. The spray voltage and capillary temperature was set to 2.3 kV and 275°C respectively. Full-scan MS spectra with a mass range of 375–1,200 m/z were acquired in profile mode using a resolution of 70,000 (maximum fill time of 250 msec or a maximum of 3e^6^ ions (automatic gain control, AGC)). Fragmentation was triggered for the top 10 peaks with charge 2–4 on the MS scan (data-dependent acquisition) with a 30-sec dynamic exclusion window (normalized collision energy was 30), and MS/MS spectra were acquired in profile mode with a resolution of 35,000 (maximum fill time of 120 msec or an AGC target of 2e^5^ ions).

##### MS protein identification and quantification

The MS data was processed as previously described (Sridharan et al., 2019). Briefly, the raw MS data was processed with isobarQuant (and identification of peptides and proteins was performed with Mascot 2.4 (Matrix Science) against a database containing *Homo sapien* Uniprot FASTA (proteome ID: UP000005640, downloaded on 14 May 2016) (Breitwieser et al., 2011) and in-house sequenced and assembled encoding sequences of Mumps Enders strain (single nucleotide variations in Table S1) along with known contaminants and the reverse protein sequences (search parameters: trypsin; missed cleavages 3; peptide tolerance 10 ppm; MS/MS tolerance 0.02 Da; fixed modifications included carbamidomethyl on cysteines and TMT10 or TMT16 plex on lysine; variable modifications included acetylation of protein N-terminus, methionine oxidation and TMT10 or 16plex on peptide N-termini). The whole cell proteome datasets were also searched with phosphorylation on S|T|Y as a variable modification.

##### MS Data analysis

All MS data analysis was performed using R (v.3.6.1).

###### Whole cell proteome data analysis

The distributions of signal sum intensities from all TMT channels was normalized with vsn (Huber et al., 2002) to correct for slight differences in protein amounts. Differential analysis was performed on log2 transformed signal sums of different stress time points and control condition using limma (Ritchie et al., 2015). Proteins with |log_2_(fold change) | > 0.5 and adjusted p-value (Benjamini Hochberg) < 0.01 were considered significantly changed.

###### Solubility proteome profiling data analysis

Data normalization for solubility proteome profiling experiments were performed based on a subset of proteins that are predominantly soluble. The NP40/SDS ratio of proteins was calculated using raw signal sum intensities. Proteins with NP40/SDS ratio between 0.8 and 1.2 represented the soluble subset. This subset was utilized for calculating the calibration and transformation parameters for vsn (Huber et al., 2002). These parameters were then applied to all proteins identified to correct for technical variations. The NP40/SDS ratio was calculated using normalized signal sum intensities and log2 transformed for performing differential analysis using limma (Ritchie et al., 2015). Proteins with |log_2_(fold change) | > 0.5 and adjusted p-value (Benjamini-Hochberg) < 0.1 were considered significantly changed.

###### Gene ontology over representation analysis

Differentially expressed human proteins from 30 µM arsenite treatment dataset were used for GO term “Biological processes” overrepresentation analysis using clusterProfiler (R Bioconductor) (Yu et al., 2012). All identified proteins in this dataset were used as the background. Standard settings were used for representing enriched GO terms (p-value cutoff: 0.05, Benjamini-Hochberg procedure for multiple testing adjustment and q-value cutoff of 0.2).

##### Nucleocapsids isolation and negative staining electron microscopy

HeLa-MuV cells were seeded on 20-cm dishes one day prior to stress treatment. After replacing the culture medium with that containing 0.5 mM As(III) and incubating the cells for ∼1 h, cells were scraped off dishes with 2 ml medium per dish, pelleted at 850 *g* for 3 min and frozen and stored at −80 °C. On the day of experiment, the cell pellets were thawed on ice and resuspend with 250 µl of lysis buffer (50 mM Tris, pH 7.5, 100 mM KAc, 2 mM Mg(Ac)_2_, 0.5 mM DTT, 0.5% IGEPAL CA-630, EDTA-free protease inhibitor cocktail, 1 U/µl Ribonuclease inhibitor Sigma) per vial of pellet. The following operations were done on ice unless specified. Cells were lysed with a syringe with a 27G ¾ needle for 5 passages. The lysate was centrifuged at 500*g*, 4 °C, for 5 min to pellet cell debris. Next, the supernatant was centrifuged at 13,000*g*, 4 °C, for 20 min. The pellet was sufficient for enriching nucleocapsids without much significant cellular contaminants. Pellets (resuspended with the lysis buffer without IGEPAL CA-630) and supernatants of each centrifugation step was used for negative staining electron microscopy (EM) experiments. Nucleocapsids in the 13,000*g* pellet resuspension were additionally supplied with heparin solution at 50 µg/ml for 2 h.

Negative staining was done with 1% uranyl acetate solution. For each sample, 2.5 µl of sample solution (1:4 to 1:100 diluted depending on concentration) was deposited on an EM copper grid (400-mesh, Plano G2400C, coated with 6 nm thick carbon produced with a Leica EM ACE600 sputter coater), incubated for 30-60 sec and manually blotted with filter paper. 2.5 µl of uranyl solution was then immediately applied to the grid and blotted away, and this step was repeated for three times. Grids were left to dry and then imaged with a Tecnai T12 EM operated at 120 kV (Thermo Fisher Scientific) at 13,000× magnification for screening and 49,000× magnification for image acquisition.

##### Sucrose gradient fractionation and MS profiling

Sucrose gradient fractionation protocol was optimized in three experiments. In the initial experiment, the cell lysate after removal of cell debris at 850*g* was first pelleted with an Optima MAX-XP Ultracentrifuge (with TLA-100 rotor, Beckman Coulter) at 35,000 rpm (47,265*g*) 4 °C, for 40 min in order to collect most nucleocapsids. The pellet was then resuspended in lysis buffer without IGEPAL CA-630, and loaded on a 5 mL 10-65% (w/v) sucrose gradient prepared with a BioComp Gradient Master instrument. Ultracentrifugation was carried out at 35,000 rpm (116,140*g*), 4 °C for 2 h in an Optima L-100XP ultracentrifuge (with SW 55 Ti rotor, Beckman Coulter). Fractionation was done manually by collecting every 200 µl volume from the top till the bottom of each centrifuge tube. Every other fraction was imaged by negative stain EM. Based on the images, the pellet at 35,000 rpm contains too much cellular material. Thus, the experiment was optimized for two more times, using the 13000*g* pellet as the starting material for a sucrose gradient of 20-70%, or 20-60% (the latter used for MS profiling analysis). Sucrose in fractions were removed by dialysis with the same lysis buffer without IGEPAL CA-630, in order to reduce background for negative staining EM. Fractions around the one showing the most abundant nucleocapsids in negative staining EM were subjected to mass spectrometry analysis as described above.

In order to estimate the relative protein amounts in the arsenate treated and control samples prior to sucrose fractionation we analyzed aliquots of the unfractionated samples from both conditions. The ratio of signal sum intensities of proteins between the unfractionated arsenate treated and control samples was calculated to correct for variations in input. The relative fold-changes of proteins in individual fractions of sucrose gradient (in control and arsenate treated sample) was calculated with reference to fraction 10. The apex of the peak for protein N, L, P and M was detected at fraction 14 in arsenate treatment fractionation and fraction 15 in control fractionation. The ratio of above-mentioned relative fold change in arsenate (fraction 14) and control (fraction 15) was calculated for N, L and P. This ratio was further corrected for variation in the input used for fractionation. The corrected ratio represented the amount of N, L and P proteins at the apex of the sucrose gradient fractionation in arsenate treated compared to control cells.

##### Plunge freezing

The isolated nucleocapsids were prepared from the lysate of HeLa-MuV cells after 1 h of 0.5 mM As(III) stress as above (13,000*g* pellet, 50 µg/ml heparin treated). Plunge freezing was done with a Vitrobot Mark IV (Thermo Fisher Scientific) set to 22 °C, 100 % humidity. Glow-discharged holey Quantifoil grids with an additional layer of continuous carbon (R2/1 + 2 nm C, Cu 200 mesh grid, Quantifoil Micro Tools) were used. 3 μl of nucleocapsids suspension was mixed with 1.5 μl of Protein-A, 10-nm colloidal gold (in the same buffer as nucleocapsids; Electron Microscopy Sciences) and deposited onto grids. Grids were blotted from both sides with blot force 0 for 2 sec and drained for 2 sec before being plunged into liquid ethane at liquid nitrogen temperature. The frozen grids were stored in sealed boxes in liquid nitrogen until imaging.

##### Sample preparation and vitrification for cellular cryo-ET

For plunge freezing of HeLa-MuV cells under different stress conditions as described above, either a Vitrobot Mark 4 or Leica EM GP (Leica Microsystems) were used. Gold Quantifoil grids (R1/4, Au 200 mesh grid, SiO2, Quantifoil Micro Tools) were glow-discharged and UV irradiated for 30 min for sterilization before being immersed in cell culture medium in 35-mm IBIDI μ-Dish. Next, HeLa-MuV cells were seeded in such dishes each containing 5-6 grids and cultured in an incubator overnight at 37 °C and 5% CO_2_. Cells cultivated on grids were plunge-frozen in liquid ethane/propane mixture at close to liquid nitrogen. The blotting conditions for the Vitrobot were set to 37 °C, 90% humidity, blot force 10, 10 sec blot time and 2 sec drain time and grids were blotted from the reverse side with the aid of a Teflon sheet from the front side. The blotting conditions for the Leica EM GP were set to 37 °C, 90% humidity, blot volume 3 μl, 2-3 s blot time and grids were also blotted from the reverse side. For grids that were used in subsequent correlative imaging, 2 μl of 1-μm crimson beads (FluoSpheres carboxylate-modified microspheres, 625/645, Thermo Fisher Scientific) diluted 1:40 from original stock were added to the grid surface from one side before blotting. Grids were stored in liquid nitrogen until usage.

##### Cryo-fluorescence light microscopy

Frozen grids were fixed into custom-made AutoGrid specimen cartridges modified for FIB preparation under shallow angles (Rigort et al., 2010) and imaged by a prototype Leica cryo-fluorescence light microscope (FLM) based on Leica TCS SP8 CFS equipped with a cryo-stage operated at liquid nitrogen temperature (Allegretti et al., 2020). The microscope includes a widefield light path and a confocal path with independent light sources and is equipped with a 50x/0.9 NA cryo-objective. The grid overview was first acquired using widefield to find focal planes. Intact grid squares with signal of interest were then imaged with the confocal path by exciting with a 552-nm laser and simultaneously detecting at 569-633 nm and 695-700 nm with two HyD detectors. After imaging, grids were stored in liquid nitrogen for the next step.

##### Cryo-focused ion beam milling

Plunge-frozen grids fixed into custom-made AutoGrids were mounted into a shuttle and transferred into an Aquilos cryo-focused ion beam/scanning electron microscope (FIB/SEM dual-beam microscope, Thermo Scientific). During FIB operation, samples were kept at constant liquid nitrogen temperature. To improve sample conductivity and reduce curtaining artifacts during FIB milling, the samples were first sputter-coated with platinum and then coated with organometallic platinum using the gas injection system (Schaffer et al., 2017). Lamellae were prepared using Gallium ion beam at 30 kV and stage tilt angles of 15°-20°. The MAPS software installed on the Aquilos microscope was used to refine eucentricity and record coordinates to prepare lamellae. Lamella preparation was conducted in a stepwise manner: rough milling with currents of 1 nA, gradually reduced to lower currents, down to 50 pA for the final polishing step (Mahamid et al., 2016). Progress of the milling process was monitored using the SEM operated at 10 kV and 50 pA. For improving conductivity of the final lamella, we sputter-coated the grid again after cryo-FIB preparation with platinum. The thickness of the platinum layer and entire lamellae was determined from the tomographic reconstructions to be approximately 5 nm and 120-250 nm, respectively.

For lamella preparation following a 3D correlative workflow, the MAPS software provides 3-point correlation (based on features of grid squares) between the cryo-FLM and cryo-SEM images for 2D navigation in order to find squares for which the cryo-FLM data were acquired. About 10 microbeads were picked in the squares of interest and correlated between the cryo-FLM stacks (3D Gaussian plot to optimize geometry center) and the SEM images, as well as the FIB images (2D Gaussian plot to optimize geometry center) using 3DCT software (v2.2.2) (Arnold et al., 2016). After choosing the signal of interest in the confocal stacks, 3DCT provided the position as the center to place the parallel rectangular milling patterns above and below the center along the Y axis on FIB images, in order to retain the signal (region of interest) in the lamella. Milling is then performed as described above. After lamella generation, 3DCT was again used to correlate the cryo-FLM data with the SEM and FIB images to confirm that the signal of interest was retained in the lamella.

##### Cryo-electron tomography data collection

Cryo-electron tomography (ET) data of isolated nucleocapsids were collected on a Titan Krios microscope operated at 300 kV (Thermo Fisher Scientific) equipped with a field-emission gun, a Quantum post-column energy filter (Gatan) and a K2 direct detector camera (Gatan). Data were recorded in dose-fractionation mode using acquisition procedures in SerialEM software (Mastronarde, 2005) (v3.7.2). Prior to the acquisition of tilt-series, montages of the grid squares were acquired at 3.106 nm/pixel to identify filamentous structures which were then confirmed as nucleocapsids at higher magnification. Tilt-series were collected for nucleocapsids using a dose symmetric scheme (Hagen et al., 2017) in nano-probe mode, calibrated pixel size at the specimen level of 1.6938 Å, defocus range 2.5 to 3.5 µm, tilt increment 3° with constant dose of 2.5 e^−^/Å^2^ for all tilts, tilt range −60° to 60° starting from 0°, a total dose of ∼102.5 e^−^/Å^2^. In total, twenty tomograms were acquired for the isolated nucleocapsids.

Cryo-ET data of FIB lamellae from cells at ∼1 h of 0.5 mM As(III) stress for subtomogram averaging purpose were collected with the same Titan Krios microscope, except the pixel size used for lamella overviews of 2.284 nm/pixel and for tilt series of 3.3702 Å, defocus at 3.25 to 3.5 µm, increment 2° with constant dose of 2.5 e^−^/Å^2^ for all tilts, tilt range −60° to 54° starting from −12° (lamella pre-tilt angle), a total dose of ∼145.0 e^−^/Å^2^. The AutoGrids with lamellae were carefully loaded with the lamella orientation (indent direction on custom-made AutoGrids) perpendicular to the tilt axis of the microscope for tilt series acquisition. For lamellae that were prepared following a 3D correlative approach, TEM overviews of lamella were overlaid with their SEM image that were already superimposed with the respective confocal oblique slice images (computed with MATLAB script), with the SerialEM registration points strategy. Two tomograms containing nucleocapsids were used for subsequent analyses.

Cryo-ET data of FIB lamellae from cells at ∼2 h of 0.5 mM As(III), ∼6 h of 30 µM As(III), or ∼6 h of 1 mM As(V) stress conditions were collected on a different Titan Krios microscope equipped with a K3 direct detector camera (Gatan). Differences in settings are pixel size for lamella overview of 2.804 nm/pixel and for tilt series of 1.631 Å, defocus at 1.75 to 3.25 µm, increment 2° with constant dose of 2.6 e^−^/Å^2^ for all tilts, tilt range −60° to 60° (starting angle depending on pre-tilt angles of lamellae), a total dose of ∼158.6 e^−^/Å^2^. In total, three tomograms containing nucleocapsids for ∼2 h of 0.5 mM As(III), one tomogram for ∼6 h of 30 µM As(III), and five tomograms for ∼6 h of 1 mM As(V) were used for subsequent analyses.

In addition to tomograms described above, which were used for subtomogram averaging, cryo-ET data of nucleocapsids *in situ* at control and ∼1 h of 0.5 mM As(III) stress were also collected with a Volta phase plate (Danev et al., 2014) (VPP, Thermo Fisher Scientific) for the purpose of better visualization. The operation of the VPP were carried out as described previously, applying a beam tilt of 10 mrad for autofocusing (Fukuda et al., 2015). Pixel size for lamella map at 2.284 nm/pixel and for tilt series at 3.3702 Å, defocus at 2 to 3.75 µm, increment 2° with constant dose of 2.20 e^−^/Å^2^ for all tilts, tilt range −64° to 50° starting from −8°, and a total dose of ∼127.6 e^−^/Å^2^ were used. In total, two tomograms containing nucleocapsids for control condition and one tomogram for ∼1 h of 0.5 mM As(III) stress were used here for visualization.

##### Tomogram reconstruction

For processing of cryo-ET data of the isolated nucleocapsids, frames of the projection movies were imported to Warp software (Tegunov and Cramer, 2019) (v1.0.9), for gain reference and beam-induced motion correction, as well as contrast transfer function (CTF) and astigmatism estimation. Prior to tomogram reconstruction in Warp, per-tilt averaged images were first imported to IMOD (Kremer et al., 1996) (v 4.9.4) for tilt series alignment. Alignment of tilt-series images was performed with patch-tracking due to insufficient gold fiducials for tracking. Final alignment was done using the linear interpolation option in IMOD. Aligned images were initially 4 times binned to a pixel size of 6.7752 Å. Reconstruction of tilt series images was done in Warp at the same pixel size by importing the transformation files from IMOD to Warp. Deconvolved tomograms at the same binning were also reconstructed in Warp for segmentation purpose.

For processing of cryo-ET data of nucleocapsids *in situ* at ∼1 h of 0.5 mM As(III) stress which were used for subtomogram averaging, the same procedures were followed except that initial tomograms were reconstructed by 4 times binning to a pixel size of 13.4808 Å. Neural network-based denoising was done in Warp at the same binning to generate tomograms for segmentation purpose.

For processing of cryo-ET data of nucleocapsids *in situ* at ∼6 h of 30 µM As(III) stress or ∼6 h of 1 mM As(V) stress which were used for subtomogram averaging, the same procedures were followed except that gain reference correction and motion correction were done in the SerialEM plugin, prior to Warp processing which started from per-tilt averaged images. Initial tomograms were reconstructed by 8 times binning to a pixel size of 13.048 Å. Neural network-based denoising was done in Warp at the same binning to generate tomograms for segmentation purpose.

For processing of cryo-ET data of nucleocapsids *in situ* at control and ∼1 h of 0.5 mM As(III) stress for visualization purpose, frames of the projection movies were gain reference corrected and motion corrected in the SerialEM plugin. Tilt-series alignment and tomographic reconstructions were done in IMOD. Alignment of tilt-series images was performed with patch-tracking. Final alignment of the tilt-series images was performed using the linear interpolation option in IMOD without contrast transfer function correction. Aligned images were 4 times binned to a pixel size of 13.4808 Å. For tomographic reconstruction by back-projection, the radial filter options were left at their default values (cut off, 0.35; fall off, 0.05).

##### Filament tracing, volume fraction and persistence length analysis

Segmentation of *in situ* nucleocapsids was performed with the Fiber Tracing module in Amira software (v6.7 and v2019.4; Thermo Fisher Scientific), using the deconvolved or denoised tomograms. The approximate outer and inner diameters, helical pitch of nucleocapsids were measured from raw tomograms by averaging the values of multiple measurements, and used for as input for the tracing parameters. The coordinates of the traced filaments were resampled with a MATLAB script to obtain equidistant points along the filament every 5.4 nm (4 pixels in datasets of 13.4808 Å/pixel). The approximate outer diameter of nucleocapsids as 21.30 nm was used for calculating the volumes of nucleocapsids per tomogram with the total lengths of nucleocapsid filaments calculated from tracing. Volume of regions per tomogram that contain nucleocapsids was obtained by manually segmenting areas where filaments were seen in Amira. The volume faction of nucleocapsids was then derived by dividing the volume of nucleocapsids to the total volume of regions containing nucleocapsids for individual tomograms. Tomograms with too few nucleocapsids (very small fractions of viral factories captured) were not used for volume fraction analysis.

Quantitative assessment of the morphology of nucleocapsid filaments was obtained by measurement of the persistence length (L_p_), the distance for which a filament’s direction persists before changing its course (Nagashima and Asakura, 1980). To determine its magnitude, we measured the difference between an angle θ_l_ between the tangents at points of distance l along the curved filament with respect to a fixed point on the filament l_0_ with angle θ_0_. Doing so for an ensemble of filaments in the tomographic volume, the correlation is determined by <cos(θ_0_-θ_l_) >= e ^−l/Lp^. Mean length of filaments for each tomogram was determined to define the length range used for fitting of the slope. Persistence lengths of microtubules, actin, and lamina were determined *in situ* to be 12.07×10^3^, 2.79×10^3^, and 555.57 nm respectively in a previous report(Mahamid et al., 2016). Nucleocapsids in our data are very flexible based on the persistence length comparison with these cytoskeletal filaments. Tomograms with too few nucleocapsids (very small fractions of viral factories captured) were not used for persistence length analysis.

##### Tomogram segmentation

For visualisation of ribosomes and membranes in the tomogram in Figure 5B, 3D convolutional neural networks for localising ribosomes and membranes are pretrained with large datasets and here used for prediction (work by de Teresa et al. in progress, https://github.com/irenedet/3d-unet/). The prediction output was inspected and cleaned manually with Amira and the final figure was generated with ChimeraX (Pettersen et al., 2021) (v1.1.1).

##### Subtomogram averaging of isolated nucleocapsids

For subtomogram averaging of the isolated nucleocapsids, nucleocapsids were first traced in Dynamo package (Castano-Diez et al., 2012) (v1.1.401) with the 4 times binned tomograms. In the Dynamo model generation interface, extremal points of relatively straight segments along each nucleocapsid were manually selected to define the central line “backbone”. Bent regions were discarded. Directionality of individual nucleocapsids were visually inspected and considered when selecting extremal points. Successive orthogonal sections along the nucleocapsid backbone were then generated such that central points of these sections were manually adjusted to define series of ordered points composing the refined backbone. After that, the Dynamo filament model with torsion type was used for generation of subtomograms (called “subunits” in Dynamo). The torsion effect in this model refers to the fact that the x direction of two subsequent subtomograms can be chosen to vary a fixed angle. Two parameters subunits_dz and subunits_dphi were set as 8 pixels (5.4 nm) and 30°, considering the helical pitch and nucleoprotein subunits per turn roughly measured/counted from tomograms, as well as for reducing the effect of missing wedge in subtomogram averaging (Wan and Briggs, 2016). Subtomograms (4 times binned, box size of 64^3^ pixels) were cropped using Dynamo with the model-defined positions and initial Euler angles were generated relative to filament axes with extra in-plane rotations derived from torsion angles (Figure S5E).

Reference-free subtomogram averaging/refinement was performed using Dynamo, adapted TOM (Nickell et al., 2005) and AV3 (Forster and Hegerl, 2007) (TOM/AV3) software toolboxes and derived MATLAB scripts, RELION (Zivanov et al., 2018) (v3.0 and v3.1), Warp and M (Tegunov et al., 2021) (v1.0.9). Initial reference was generated by averaging subtomograms (4 times binned) from a single long nucleocapsid without alignment. A cylindrical mask was generated in Dynamo according to the initial average, and used as alignment mask. Other masks were left as default. Subtomograms from the single long nucleocapsid were iteratively aligned against the initial reference in Dynamo, performing Euler angles and Cartesian shifts search. The resulting average of the single long nucleocapsid was used to align all subtomograms extracted from 20 tomograms, with the first two Euler angles restrained to not allow flipping during angular search. After the first round of rough alignment with a low-pass filter of ∼35 Å, the resulting average showed polarity. To determine the directionality of individual nucleocapsid, averages of subtomograms belonging to individual nucleocapsid with refined positions and angles were visually inspected. Subtomograms of nucleocapsids with opposite directionality to the initial long nucleocapsid were flipped by modifying their Euler angles. After directionality check, rounds of iterative alignment were done until convergence. The refined coordinates of subtomograms were used for reconstructing 2 times binned subtomograms in Warp.

The subtomograms were then split into odd and even sets at 2 times binning for alignment using TOM/AV3 toolboxes. The odd and even sets were aligned independently using the average of each set as initial reference. Low-pass filter and angular search range/sampling parameters were adjusted based on the Fourier Shell correlation (FSC) plots after each round of alignment until convergence. After that, distance-based cleanup of subtomograms was done with MATLAB scripts to remove particles that were too close due to shift during alignment and the ones with higher constrained cross correlation (CCC) values were kept. The refined coordinates of subtomograms were used for reconstructing unbinned subtomograms (box size of 256^3^) with per-particle CTF models in Warp.

Unbinned subtomograms were averaged with refined Euler angles from TOM/AV3 alignment and intensity-inverted to generate initial reference for RELION 3D auto-refinement. Then, standard 3D auto-refinement was performed without symmetry, with a soft-edged sphere-multiplied cylindrical mask which was generated using relion_helix_toolbox (outer diameter 220 Å, sphere percentage 0.55). A 30 Å lowpass-filtered map generated from TOM/AV3 alignment was used as an initial reference. 1,209 subtomograms led to a 10 Å map with no symmetry applied. Helical parameters were estimated with relion_helix_toolbox, resulting in an average helical rise of 4.46 Å and helical twist of −27.08° (left-handed, ∼13.3 subunits per turn). Next, helical symmetry was imposed in RELION 3D auto-refinement with local symmetry search, resulting in the refined average helical rise of 4.16 Å and helical twist of −27.16°. After RELION classification with helical symmetry imposed, two classes of distinct helical parameters were separated: the majority class (77.7%, 4.21Å/-27.17°, map at 6.5 Å after auto-refinement) and the minority class (21.3%, 3.51Å/-26.9°, map at 7.3 Å after auto-refinement).

Positions and orientations of RELION symmetry-expanded subtomograms of the majority class were next used for refining tilt series alignment and CTF models in M software (Tegunov et al., 2021). After resolution convergence was reached in M (4.8 Å), new non-symmetry expanded subtomograms were reconstructed in Warp. One more round of RELION 3D auto-refinement imposing helical symmetry (which remained the same after M refinement) resulted in a final map of 4.5 Å for the majority class within the reconstruction mask. The minority class was refined in a newer version of M which allowed to set helical symmetry parameters. After resolution convergence was reached in M, subtomograms were again reconstructed in Warp. One more round of RELION 3D auto-refinement imposing helical symmetry (remaining the same after M refinement) resulted in a final map of 6.3 Å for the minority class within the reconstruction mask with the 0.143 criterion.

Maps used for structural analysis were obtained by filtering to the estimated resolution using the RELION Post-processing module with the same sphere-multiplied cylindrical mask. Visualization of density maps were done with UCSF Chimera (Pettersen et al., 2004) (v1.13.1)/ChimeraX. FSC plots were obtained with the RELION Post-processing module using the same sphere-multiplied cylindrical mask. Local resolution estimation was also done in RELION with the Local resolution module providing the B-factors obtained from post-processing jobs and the masks. Local resolution maps were rendered with Chimera.

##### Subtomogram averaging of straight nucleocapsids *in situ*

For subtomogram averaging of the relatively straight nucleocapsids *in situ* (at ∼6 h of 30 µM As(III) and ∼6 h of 1 mM As(V) stress), the coordinates of the traced filaments were resampled with a MATLAB script to obtain equidistant points along the filament at every 5.2 nm (4 pixels of 8 times binned data, 13.4808 Å/pixel) and with a torsion angle (30°) assigned in similar way as for the isolated nucleocapsids (Figure S7D). These equidistant points were used as center points for extraction of subtomograms (box size of 32^3^) with Dynamo crop function. Alignment of 8 times binned subtomograms were done with Dynamo. Alignment started with the longest nucleocapsid in one tomogram exhibiting the longest persistence length. As described above, initial average was generated and used as reference for alignment of subtomograms in this longest nucleocapsid. The resulting average was used for alignment of subtomograms from four long nucleocapsids in this tomogram. To determine the directionality of the other three nucleocapsids, averages of subtomograms belonging to individual nucleocapsid with refined positions and angles were visually inspected. Subtomograms of nucleocapsids with opposite directionality to the longest nucleocapsid were flipped by modifying their Euler angles. After directionality check, alignment was re-done with subtomograms from these four nucleocapsids. The resulting average was next used to align all subtomograms in six tomograms for one round of angular and translational search. Directionality check was done for individual nucleocapsid as described above. Only nucleocapsids showing clear directionality amongst those longer than 100 nm were kept for further processing. Subtomograms of nucleocapsids with opposite directionality to the longest nucleocapsid were flipped by modifying their Euler angles. Next, two more rounds of iterative alignment were done until convergence. Similarly, distance- and CCC-based cleaning of subtomograms was done with MATLAB scripts. The refined coordinates of subtomograms were used for reconstructing 4 times binned subtomograms in Warp.

The 4 times binned subtomograms were then aligned using TOM/AV3 toolboxes. After three rounds of iterative alignment, distance- and CCC-based cleanup of subtomograms was done. The refined coordinates of subtomograms were used for reconstructing two times binned subtomograms (box size of 128^3^) with per-particle CTF models in Warp.

The 2 times binned subtomograms were used for RELION 3D auto-refinement, following similar procedures as described for the isolated nucleocapsids. 2,178 subtomograms led to a 22 Å map with no symmetry applied. Helical parameters were estimated with relion_helix_toolbox and helical symmetry was imposed in RELION 3D auto-refinement with local symmetry search, resulting in the refined average helical rise of 4.24 Å and helical twist of −27.19°, and improved map resolution to 8.9 Å. RELION classification with helical symmetry imposed resulted in classes with very similar helical parameters, and were thus combined for further refinement. Positions and orientations of RELION helical symmetry-expanded subtomograms were next used for refining tilt series alignment and CTF models in M software. After resolution convergence was reached in M at 2 times binning, new non-symmetry expanded subtomograms were reconstructed in Warp. One more round of RELION 3D auto-refinement imposing helical symmetry resulted in a map of improved resolution at 7.2 Å and of the same helical parameters as the majority class in subtomogram averaging of isolated nucleocapsids (rise as 4.21Å and twist as −27.17°).

Unbinned subtomograms (box size of 256^3^) were then reconstructed in Warp at refined positions and subjected to RELION 3D auto-refinement. This step of 3D auto-refinement did not improve the resolution and classification resulted in very minor difference in helical parameters. The map was further refined in M which allowed to set helical symmetry parameters. After resolution convergence was reached in M, subtomograms were again reconstructed in Warp. One more round of RELION 3D auto-refinement imposing helical symmetry (remaining the same after M refinement) resulted in a final map of 6.5 Å within the reconstruction mask with the 0.143 criterion.

The map used for structural analysis was obtained by filtering to the estimated resolution using the RELION Post-processing module with the sphere-multiplied cylindrical mask.

##### Subtomogram averaging of curved nucleocapsids *in situ*

For subtomogram averaging of the curved nucleocapsids *in situ* (at ∼1 h of 0.5 mM As(III) stress), the coordinates of the traced nucleocapsids (done in Amira) were over-sampled with a MATLAB script to obtain uniformly distributed positions on the surfaces of nucleocapsid at every 2.7 nm (4 pixels of 2 times binned data), with initial Euler angles assigned at each surface position, and the new Z axis assigned perpendicular to the nucleocapsid axis (Figure S7A). These surface positions were used as extraction points for subtomogram reconstruction in Warp.

Alignment of 2 times binned subtomograms (box size of 36^3^) was done with TOM/AV3 toolboxes. The initial average was generated by aligning and averaging subtomograms of the longest nucleocapsid in a tomogram acquired with VPP, which has better contrast (signal-to-noise ratio) at low spatial frequency range. The VPP-average was used for alignment of subtomograms of all nucleocapsids in the two tomograms acquired without VPP. After 5 iterations of initial alignment, nucleocapsid directionality was checked by visual inspection of subtomogram averages of individual nucleocapsid. However, determination of the directionality was difficult due to poor signal-to-noise ratio of individual average. Thus, sampling was re-done at equidistant points along the central line of nucleocapsids and subtomograms were extracted for generation of averages of individual nucleocapsid, only for determination of directionality. Nucleocapsids showing clear directionality amongst those longer than ∼130 nm were kept for further processing. Subtomograms (oversampled) of nucleocapsids with opposite directionality to the VPP-average were flipped by modifying their Euler angles. Next, the VPP-average was again used to align the remaining 27,464 subtomograms (oversampled) until convergence. After that, distance- and CCC-based cleaning of clashing subtomograms was done with MATLAB scripts, with 4 pixels (2.7 nm) as a threshold. The refined coordinates of the 10,702 remaining subtomograms were used for reconstruction of subtomograms at 2 times binning in Warp with the same box size.

The re-extracted 2 times binned subtomograms were used for RELION 3D Classification using the average converted from TOM/AV3 alignment as a reference (filtered to 40 Å). Four classes were generated and one class (2882 particles) was discarded due to poor signal to noise ratio. The other three classes (2452, 2726, 2642 particles each) were separately refined, all resulting in maps of ∼30 Å in resolution. Further refinement using smaller masks did not improve maps, but resulted in too little content to be compared with higher resolution maps of isolated nucleocapsids. The maps used for structural analysis were obtained by filtering to the estimated resolution using the RELION Post-processing module with the refinement mask.

##### Model building

To build a pseudo-atomic model for the majority class of the isolated nucleocapsids resolved to 4.5 Å, an initial model of MuV-N was generated with the homolog N protein of parainfluenza virus 5 (PIV5) (PDB: 4XJN) as a template using I-TASSER (Roy et al., 2010). The initial model was first fitted into the map of the majority class as a rigid-body, together with six RNA bases from the PIV5 model using Chimera. Only the core part of N (amino acid 3-405) was used for model building because the density for C-terminal flexible region was missing. Then, the combined model of N and RNA was subjected to one round of automated refinement using Phenix (Liebschner et al., 2019) (v 1.18-3845) with minimization_global, local_grid_search, simulated_annealing, and ADP strategies, and restraining rotamers, Ramachandran and secondary structures. Next, three copies of the refined N and RNA were generated and rigid-body docked to EM density corresponding to the subunit at N_i+1_ position with respect to the original one (N_i_). Two additional subunits were docked into the upper turn in order to take interaction interfaces into account. After that, the tetramer model was per-residue manually refined in *Coot* (Emsley et al., 2010) (v0.9) using Peptide and Ramachandran restraints, with attention on the first monomer and molecular interfaces. Regions poorly fitted were deleted and re-built. N- and C-termini were checked and more residues added in case densities were seen. The RNA bases were modeled as poly-Uracils considering the averaging effect. Density modification strategy in Phenix.ResolveCryoEM was used to generate a map for aiding model inspection in *Coot*, but automated refinement and validation were done against the original map. The refined model of the first monomer was again copied and rigid body docked into the density map for a second round of automated refinement with Refine NCS operators on. Additional rounds of manual inspection and automated refinement were performed until convergence and Refine NCS operators was not used in the last round. The real-space cross-correlation of each residue to the map was calculated and plotted in order to evaluate the quality of the model (Figure S6C). The masked FSC between the original map and a simulated map calculated from the tetramer model at 0.5 cutoff gave a resolution estimate of 4.5 Å, similar to the estimate from the half map FSC 0.143 cutoff. MolProbity statistics were computed to ensure proper stereochemistry. Monomer of the model was used for rigid-body fitting into the map of straight nucleocapsids *in situ* (6.5 Å) with Chimera.

#### QUANTIFICATION AND STATISTICAL ANALYSIS

The details of the quantification and all statistical analyses are included in figure captions or the relevant sections of METHOD DETAILS.

## Notes

### Competing Interest Statement

A.A.H. is a founder and scientific advisory board member of Dewpoint Therapeutics and a founder of Caraway Therapeutics. All authors declare no competing interests.

### Summary of Updates

Section "Stress alters protein interaction networks within viral factories and induces stabilization of the MuV replication machinery" updated to include new proteomics data; Figure 2 and figure S3 revised to include new proteomics data; all figures revised to accommodate the changes; The introduction and discussion sections expanded to provide relevant background and context on persistent viral infection, bio-molecular condensates, structures of viral nucleocapsids;

