## Supplementary material for "Molecular mechanisms of stress-induced reactivation in mumps virus condensates": Figures S1 to S7; Tables S1-S5 and Video S1-S4 captions; Tables S6, S7

### Figure S1

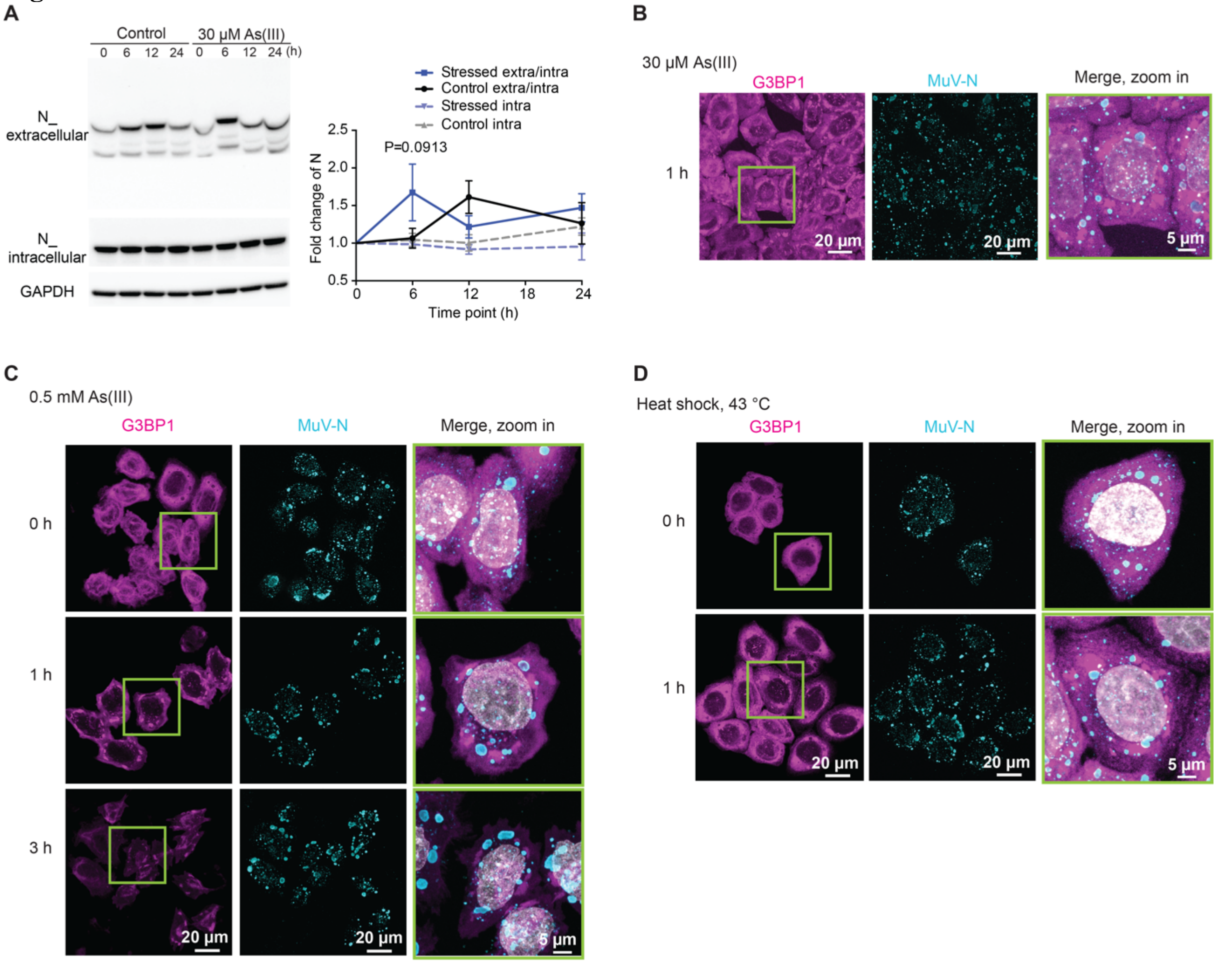


**Figure S1. Effect of different stress conditions on viral particle release and viral factories coarsening in HeLa-MuV cells, related to Figures 1 and 2**

(A) Representative western blot and quantification of MuV N levels in the cell culture medium (N_extracellular) and inside cells (N_intracellular) during 24 h of 30 μM As(III) stress or of control unstressed cell samples collected at the same time points. Glyceraldehyde 3-phosphate dehydrogenase (GAPDH) was used as a loading control. Lower bands in N_extracellular are likely degraded forms of N (Brgles et al., 2016). Ratio of MuV N_extracellular to N_intracellular (labeled as extra/intra for plots) and levels of N_intracellular (labeled as intra for plots) are normalized to levels at 0 h for each condition. Data are the mean ± SEM ($n$ = 4).

(B) MIPs of MuV factories in HeLa-MuV cells detected by anti-N immunostaining (cyan) in HeLa mCherry-G3BP1 (magenta) at 1 h of 30 μM As(III) stress, completing data presented in Figure 2B. Magenta: mCherry-G3BP1; Grey: DNA stained with DAPI.

(C) MIPs of MuV factories in HeLa-MuV cells during 3 h of 0.5 mM As(III) stress. Quantification is provided in Figure 2E.

(D) MIPs of MuV factories in HeLa-MuV cells after 1 h of heat shock (HS) at 43 °C in comparison to control. Quantification is provided in Figure 2F.

### Figure S2

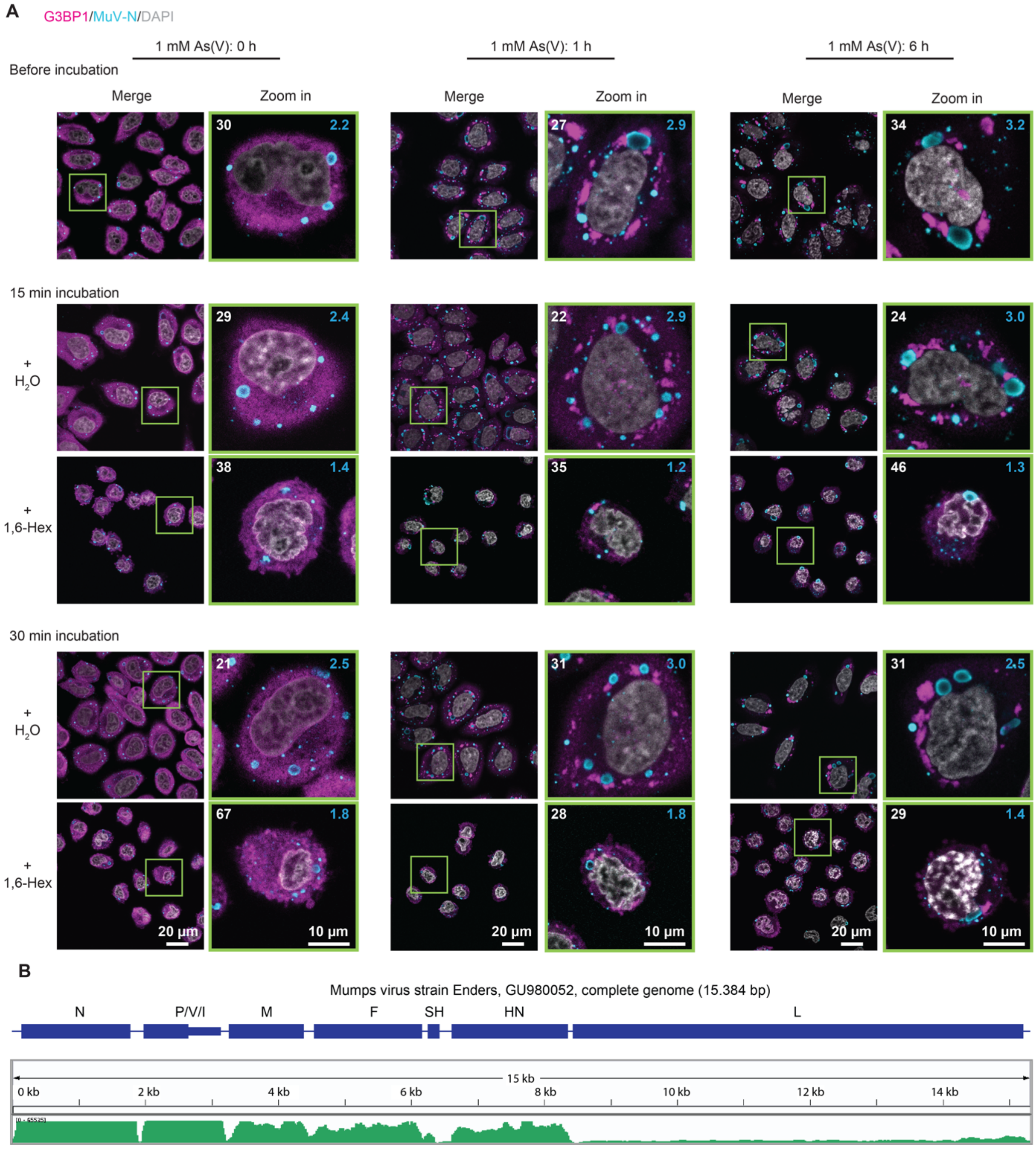


**Figure S2. Probing liquid-like properties of MuV factories and expression levels of MuV genes in HeLa-MuV cells, related to Figures 2 and 3**

(A) Exemplary image of hexanediol treatment on MuV factories in HeLa-MuV cells along stress treatment. See also Figure 2G.

(B) Transcriptome sequencing of HeLa-MuV cells after ~1 h of 0.5 mM As(III) stress. Transcripts were mapped to the MuV genome to reveal its transcription profile (green). The unit of the coverage tracks is number of reads per genomic position, scaled at 0-65535. P/V/I are co-transcriptional products; little is known about the function of I protein and it is not discussed in this study.

### Figure S3

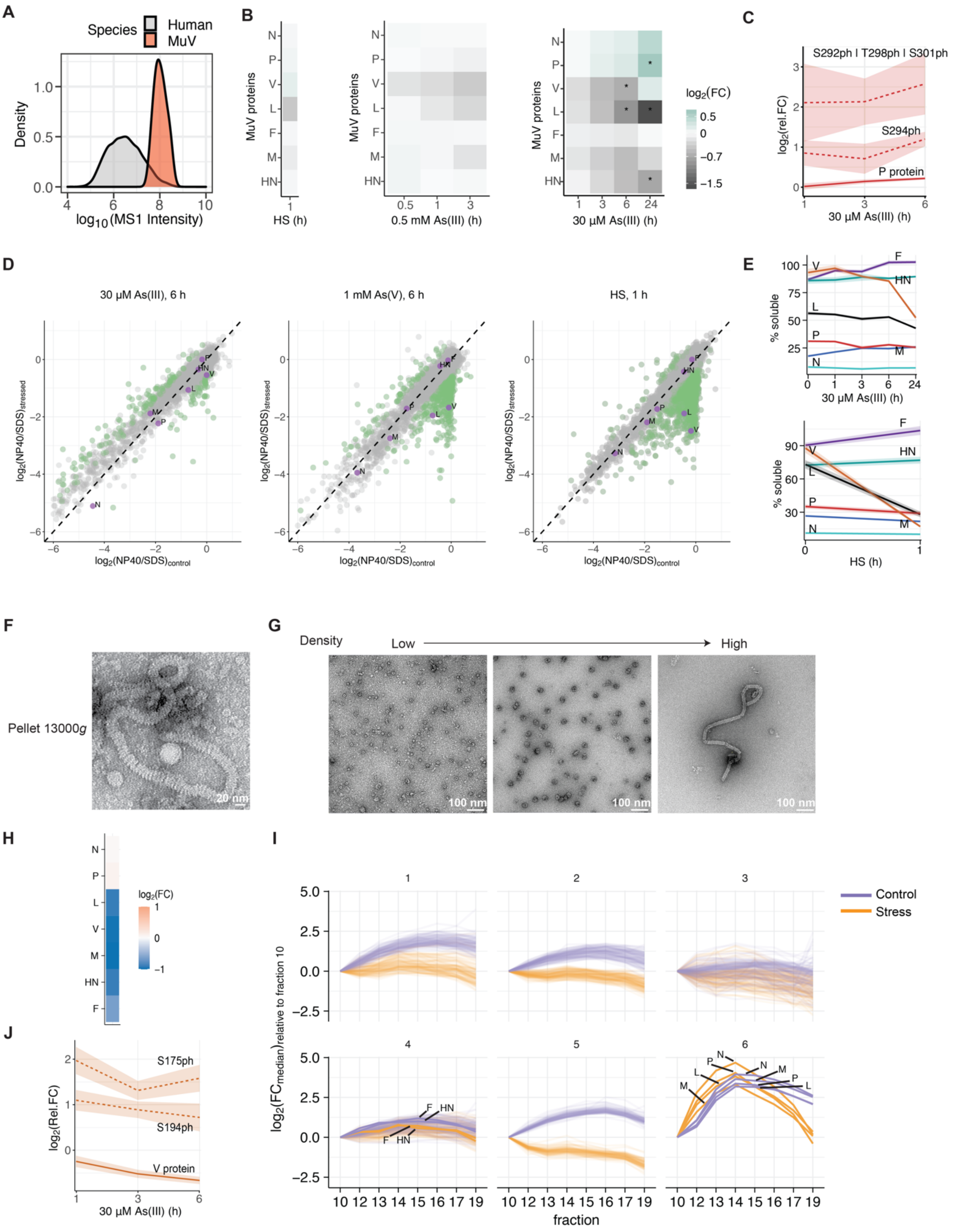


**Figure S3. Proteomics of HeLa-MuV, related to Figure 4**

(A) Density plot representing the distribution of MS1 intensity of the MuV and human proteomes in unstressed Hela-MuV cells.

(B) Heat map representation of relative fold change (FC) of all MuV proteins inside cells at indicated time points of heat shock (HS) at 43 °C, 0.5 mM As(III) or 30 μM As(III) stress, in comparison to the unstressed control (*n* = 3). Asterisks indicate proteins with $|\log_2(\text{fold change})| > 0.5$ and adjusted p-value < 0.01 (Benjamini-Hochberg method).
(C) Relative fold change of P along 30 μM As(III) stress time course relative to control: unmodified protein (solid line) and phosphopeptides (dashed lines). Lines and shaded areas are mean and SEM (*n* = 3). See also Figure 4B.
(D) Scatter plots representing the solubility of viral and host proteins in control cells (x-axis) and under the indicated stress condition (y-axis). Solubility is defined by NP40/SDS ratio, indicating the proportion of soluble protein in the lysate pelleting assay. Purple dots: viral proteins; green dots: host factors that exhibited significant change in solubility; grey dots: host factors that did not exhibit significant change in solubility. Proteins with $|\log_2(\text{fold change})| > 0.5$ and adjusted p-value (Benjamini-Hochberg method) < 0.1 were considered significantly changed. Data are mean values (*n* = 2).
(E) Solubility profiles of all viral proteins along 30 μM As(III) stress time course, or after heat shock stress, related to Figure 4D. Solubility is defined the same way as in (D). Lines and shaded areas are mean and SEM (*n* = 2).
(F) Representative negative staining transmission electron microscopy (TEM) image of nucleocapsids enriched in the 13,000*g* pellet of the lysate from HeLa-MuV cells after ~1 h of 0.5 mM As(III) stress.
(G) Representative negative staining TEM images of fractions from sucrose gradient centrifugation experiment.
(H) Heat map representation of relative fold change (in $\log_2$ scale) of all MuV proteins in 13,000*g* pellet (input of sucrose gradient fractionation) after 6 h of 1 mM As(V) stress in comparison to the unstressed control. Median values are shown (*n* = 3).
(I) Hierarchical clustering of sucrose gradient fractionation profiles (median relative fold change compared to Fraction 10, *n* = 3) of all proteins in unstressed control and 6 h of 1 mM As(V) stress. Thick lines: MuV proteins; thin lines: HeLa proteins. See also Figures 4E and 4F.
(J) Relative fold change of V along 30 μM As(III) stress time course relative to control: unmodified protein (solid line) and phosphopeptides (dashed lines). Lines and shaded areas are mean and SEM (*n* = 3). See also Figure 4G.

### Figure S4

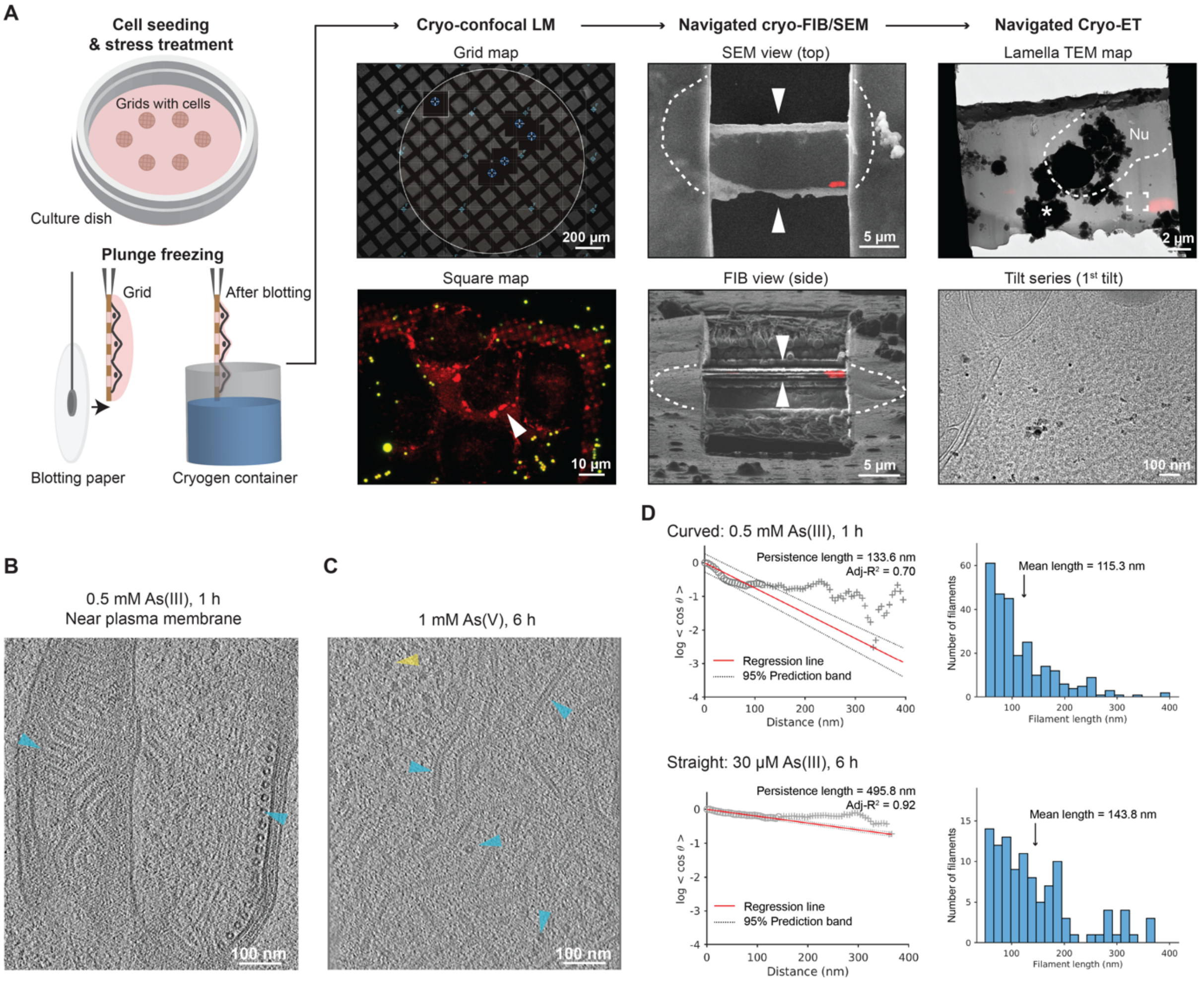


**Figure S4. Workflow of *in situ* cryo-correlative electron tomography and tomogram analysis, related to Figure 5**

(A) 3D cryo-correlative light and electron microscopy (CLEM) workflow targeting unlabeled MuV factories in vicinity of stress granules (SGs). MuV factories were frequently seen in proximity of SGs (see also Figure 2A, B). HeLa-MuV cells were seeded and cultured on TEM grids in a culture dish. At different time points of stress, grids with cells were plunge-frozen, followed by imaging with a cryo-confocal light microscope (LM). Grid squares with cells showing SG signals (red; indicated with white arrowhead) and optimal fluorescent beads (yellow) distribution in grid map overview were chosen for imaging in confocal mode. Lamellae were then prepared by cryo-FIB at SG locations, using the fluorescent beads as fiducials for 3D correlation between LM and SEM/FIB images. Correlating SEM/FIB images with 3D LM signals after lamella preparation confirmed SG location preservation (red signal) on lamella. Overlaying the SEM image with transformed fluorescent signal onto the cryo-TEM image of the lamella allowed identification of SG location and informed tilt series acquisition (indicated by frame) at nearby regions where filamentous structures were seen and later identified to be MuV nucleocapsids. Nu: nucleus. Asterisk: ice contaminants introduced during transfers between the cryo-microscopes.

(B) A 6.74 nm-thick tomographic slice of nucleocapsids aligned near the plasma membrane. Cyan arrowheads: nucleocapsids.

(C) Representative tomographic slice showing MuV factories at 1 mM As(V) stress condition. Cyan arrowheads: nucleocapsids (both top and side views, two of each); yellow arrowheads: ribosomes. Slice thickness: 6.52 nm. See also Figure 5C.

(D) Examples of persistence length measurement (left) and histogram of nucleocapsid length (right) derived from tracing of nucleocapsids in two tomograms representing curved and straight nucleocapsids, obtained at indicated stress conditions. 3D coordinates along all traced nucleocapsids within one tomogram were used to calculate the "tangent-correlation length" (apparent persistence length in the filament ensemble), which is inversely related to the curvature of filament (see also STAR Methods). Mean length of filaments per tomogram was determined to define the length range used for fitting.

Figure S5

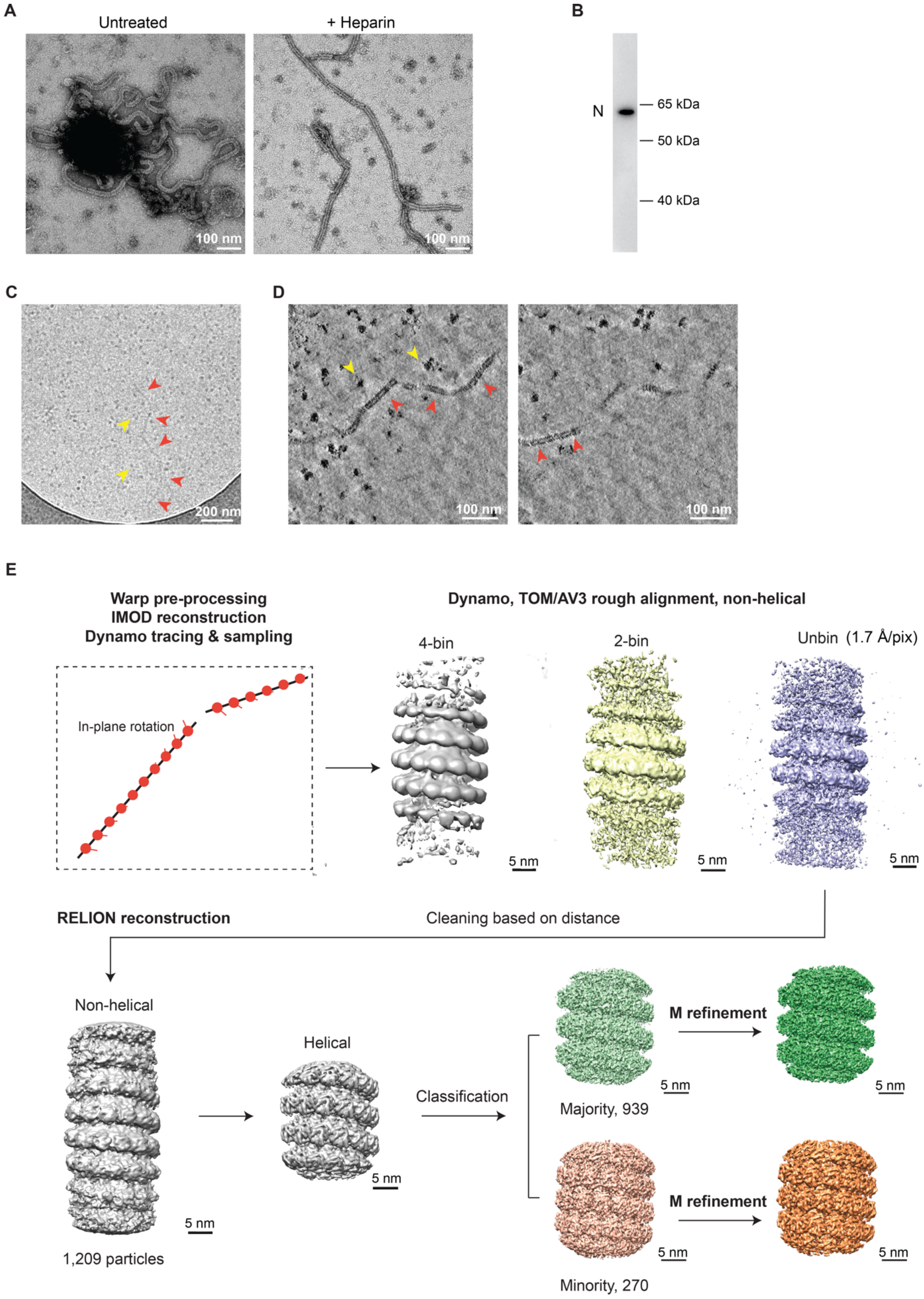


**Figure S5. Structure determination of extracted MuV nucleocapsids, related to Figure 6**

(A) Negative staining TEM of nucleocapsids enriched from cell lysates in the 13,000*g* pellet (left), and the effect of heparin addition on nucleocapsid morphology.

(B) Western blot detects full-length MuV N protein in the 13,000*g* pellet.

(C) Representative cryo-TEM image of extracted nucleocapsids (13,000*g* pellet, red arrowheads) at intermediate magnification. Nearby ribosomes are labeled (yellow arrowheads).

(D) Two 3.39 nm-thick tomographic slices of the nucleocapsid in (C), 12 nm apart in the z direction. Red arrowheads: straight segments of a long nucleocapsid; yellow arrowheads: ribosomes.

(E) Subtomogram averaging workflow for the extracted nucleocapsids from HeLa-MuV cells after ~1 h of 0.5 mM As(III) stress. Frames in each tilt (20 tilt series in total) were aligned, averaged, and CTF-estimated in Warp (CTF: contrast transfer function). Averaged tilt series images were aligned and reconstructed into tomograms in IMOD. Alignments were imported into Warp for 3D-CTF estimation, tomogram reconstruction and application of a deconvolution filter. Nucleocapsids were traced manually in Dynamo. 1209 center positions along the nucleocapsids were used to crop subtomograms with orientations assigned based on filament directions, with 30° in-plane rotation initially assigned to subsequent subtomograms. Template-free alignment was done in Dynamo at 4 times binning. Alignment results and average were used for subsequent processing of 2 times binned and unbinned subtomograms in TOM/AV3. No helical symmetry was applied at these stages. 3D refinement in RELION was first performed without applying symmetry until helical parameters could be determined in RELION. Helical reconstruction and classification were then done to separate subtomograms of different helical parameters. M refinement with symmetry expansion subsequently improved resolution and map quality. Except for the Dynamo alignment step, CTF-corrected subtomograms and 3D CTF models were all reconstructed in Warp. Overlapping particles were removed in intermediate steps and particle numbers used in each step are indicated. Also see STAR Methods.

**Figure S6**

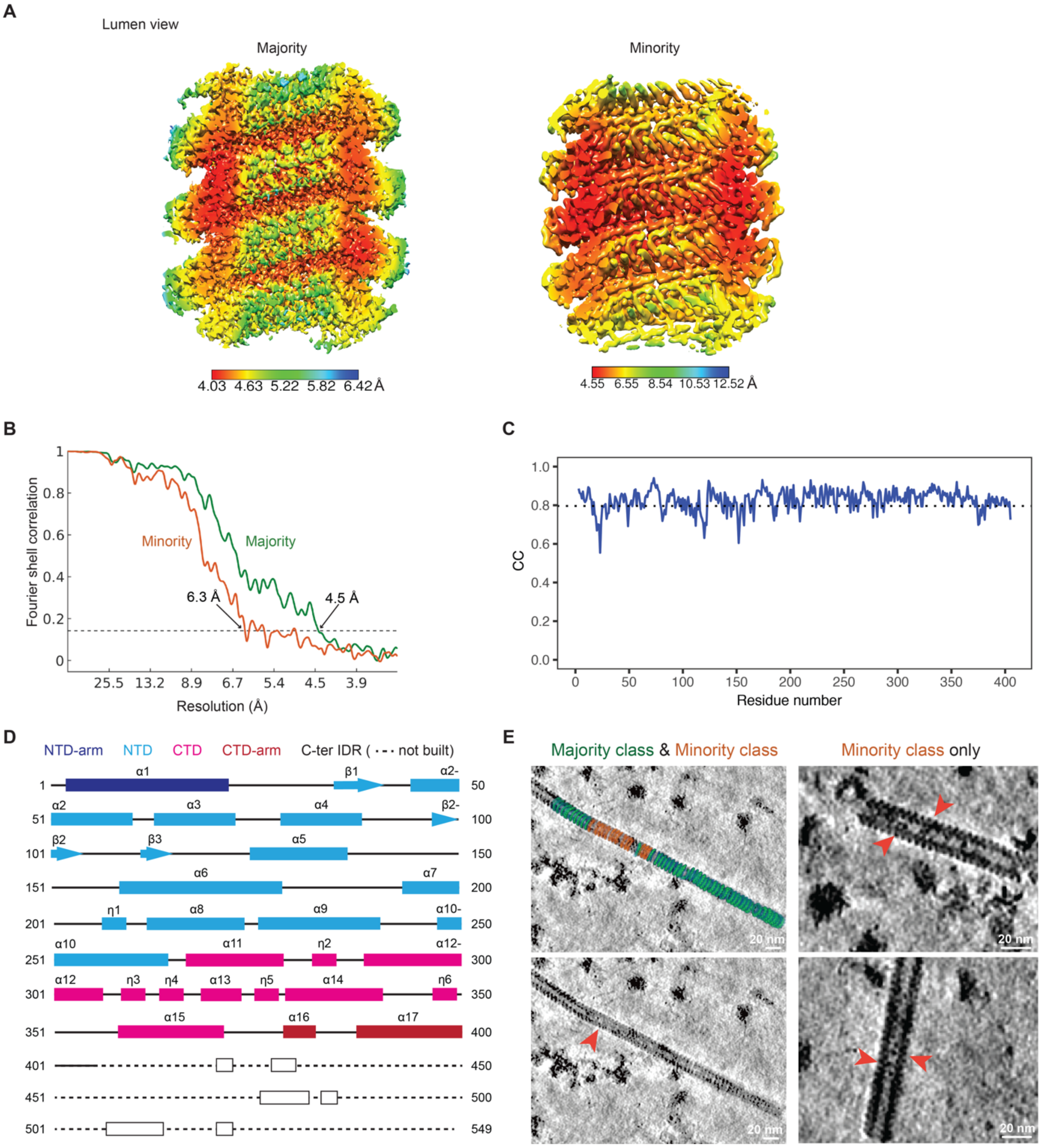


**Figure S6. Structural analysis of the two classes in extracted nucleocapsids, related to Figure 6**

(A) Local resolution maps for the majority and minority class averages.

(B) Fourier shell correlation (FSC) curves between the two independently refined half-sets for the majority and minority maps shown in (A), with the 0.143 criterion.

(C) Plot of per residue cross correlation (CC) coefficients between the generated atomic model for N (amino acid 3-405) and the segmented subunit in the majority class map. The dotted line shows the average CC at a value of 0.80.

(D) Illustration of the secondary structure of N. Secondary structure of the C-ter IDR was predicted with PSIPRED.

(E) Mapping of the majority and minority classes into tomograms. Red arrowheads indicate minority class mapped regions, which show densities at nucleocapsid lumen, possibly representing the C-ter IDR of N protein in this conformation.

**Figure S7**

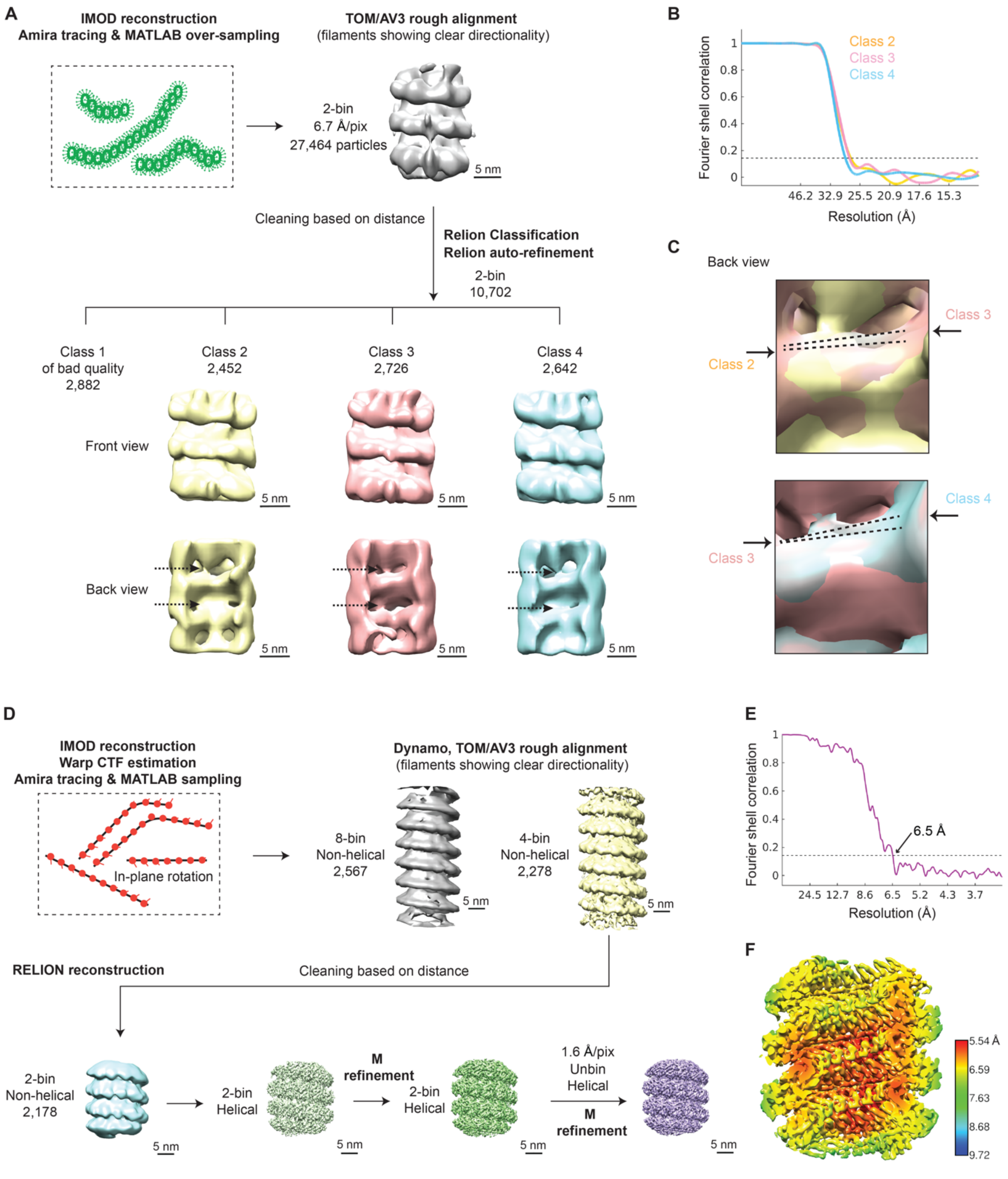


**Figure S7. Subtomogram averaging for nucleocapsids *in situ*, related to Figure 7**

(A) Subtomogram averaging workflow for the curved nucleocapsids inside HeLa-MuV cells after 1 h of 0.5 mM As(III) stress, the same stress condition used for structure determination of isolated nucleocapsids. Nucleocapsids were traced by filament tracing in Amira and sampled with custom-made MATLAB scripts. An over-sampling strategy for particles at the nucleocapsid surface was used because the filaments are too curved to be averaged by helical reconstruction. After initial alignment in TOM/AV3, RELION classification resulted in three classes with mixed helical rises. The connecting densities between turns in back view (luminal, arrows) show high heterogeneity within each average, leading to difficulty in determination of exact helical parameters. Also see STAR Methods.

(B) FSC curves for the class averages shown in (A), with the 0.143 criterion.

(C) Map comparison between class averages in (A). The dotted lines qualitatively represent the different inclination of the helical turns in the compared classes.

(D) Subtomogram averaging workflow for the straight nucleocapsids inside HeLa-MuV cells after 6 h of 30 μM As(III) stress or 1 mM As(V) stress. Particles from 6 tomograms at the two prolonged stress conditions were merged. Averaged tilt series images were CTF-estimated and reconstructed into tomograms in Warp with alignment files from IMOD, and then denoised in Warp. Nucleocapsids were traced by filament tracing in Amira and sampled with custom-made MATLAB scripts. Initial orientations of subtomograms were assigned based on filament directions with 30° in-plane rotation angles for subsequent subtomograms. Template-free alignment was done in Dynamo at 8 times binning. Alignment results and average were used for processing of 4 times binned subtomograms in TOM/AV3. No helical symmetry was applied at these stages. 3D refinement in RELION was first done at 2 times binning without applying symmetry, until helical parameters could be determined. Helical reconstruction was then done to improve the resolution. M refinement with helical symmetry subsequently improved the map quality. Helical reconstruction was finally done with unbinned particles. Classification trials resulted in classes with very similar helical parameters, and are not shown here. CTF-corrected subtomograms and 3D CTF models were all reconstructed in Warp. Overlapping particles were removed in intermediate steps, and particle numbers at each step are indicated. See also STAR Methods.

(E) FSC curve for the final map shown in (D), with the 0.143 criterion.

(F) Local resolution map for the final map shown in (D).

**Captions for Tables S1 to S5**

Tables are provided in separate text and Excel files.

**Table S1, related to Figure S2B**

Single nucleotide variations of the MuV identified in this study based on transcriptome sequencing data compared to the Enders strain sequence

**Table S2, related to Figure 4**

Mass spectrometry quantification of viral and host protein expression levels during the time course of different stress treatment

**Table S3, related to Figure 4**

Mass spectrometry quantification of viral protein phosphorylation levels during the time course of different stress treatment

**Table S4, related to Figure 4**

Mass spectrometry solubility profiling of viral and host proteins during the time course of different stress treatment

**Table S5, related to Figure 4**

Mass spectrometry profiling of sucrose gradient centrifugation fractions of nucleocapsid complex in control and 6 h of 1 mM As(V) stress conditions

| | Majority isolated (EMD-13133) (PDB 7OZR) | Minority isolated (EMD-13136) | Straight *in situ* (EMD-13137) | Curved *in situ* |
|---|---|---|---|---|
| **Data collection and processing** | | | | |
| Magnification | 81,000 | 81,000 | 53,000 | 42,000 |
| Voltage (kV) | 300 | 300 | 300 | 300 |
| Detector | Gatan K2 | Gatan K2 | Gatan K3 | Gatan K2 |
| Electron exposure ($e^{-}/Å^2$) | 102.5 | 102.5 | 158.6 | 145.0 |
| Defocus range (μm) | 2.5-3.5 | 2.5-3.5 | 1.75-3.25 | 3.25-3.5 |
| Pixel size (Å) | 1.6938 | 1.6938 | 1.631 | 3.3702 |
| Symmetry imposed | Helical | Helical | Helical | C1 |
| Final no. of segments/particles | 939 | 270 | 2,178 | 7,820 |
| Helical rise (Å) | 4.21 | 3.51 | 4.21 | NA |
| Helical twist (°) | -27.17 | -26.9 | -27.17 | NA |
| Map resolution at 0.143 FSC (Å) | 4.5 | 6.3 | 6.5 | 30 |
| **Refinement** | | | | |
| Initial model used (PDB code) | 4XJN | | | |
| Model resolution at 0.5 FSC (Å) | 4.5 | | | |
| Map sharpening *B* factor ($Å^2$) | -56.3 | | | |
| Model composition | | | | |
| Non-hydrogen atoms | 3320 | | | |
| Protein residues | 403 | | | |
| Nucleotide residues | 6 | | | |
| *B* factors ($Å^2$) | | | | |
| Protein | 86.50 | | | |
| Nucleotide | 84.56 | | | |
| R.m.s. deviations | | | | |
| Bond lengths (Å) | 0.005 | | | |
| Bond angles (°) | 0.709 | | | |
| Validation | | | | |
| MolProbity score | 2.17 | | | |
| Clashscore | 16.12 | | | |
| Poor rotamers (%) | 0.00 | | | |
| Ramachandran plot | | | | |
| Favored (%) | 92.77 | | | |
| Allowed (%) | 7.23 | | | |
| Disallowed (%) | 0.00 | | | |

**Table S6. Cryo-ET data collection, refinement and validation statistics, related to STAR Methods**

| Primer Name | Primer Sequence (5' -> 3') |
|---|---|
| **Primers used for cDNA generation** | |
| 3'-UTR genomic forward primer | AGCTTGATCCTCACCTTCACC |
| L-gene antigenomic reverse primer | CATTTGGTAACTGGCCATGC |
| RNase P_reverse primer | GAGCGGCTGTCTCCACAAGT |
| **Primers and probes against gene of interest** | |
| N-gene forward primer | GTATGACAGCGTACGACCAACCT |
| N-gene reverse primer | GCGACCTTGCT CTGGTATT |
| N-probe-LC610 | LC610_CTGGATCTGCTGATCGACGAT_BHQ1 |
| P-gene forward primer | GAGAAACCTGGAACCTCAAC |
| P-gene reverse primer | TTGGACCCAGCTGAGGCTC |
| P-probe-Cy5 | Cy5_CTGCTCAAGGCCAGACAATCCAAGAGGA_BHQ2 |
| F-gene forward primer | TCTCATCTATAGCAGGGAGTTATAT |
| F-gene reverse primer | GTTAGACTTCGACAGTTTGCAACAA |
| F-probe-FAM | 6-FAM_AGGCGATTTGTAGCACTGGATGGAACA_BHQ1 |
| RNaseP-gene forward primer | AGATTTGGACCTGCGAGCG |
| RNaseP-gene reverse primer | GAGCGGCTGTCTCCACAAGT |
| RNaseP-probe-HEX | HEX_TTCTGACCTGAAGGCTCTGCGCG_BHQ1 |

**Table S7. Primers used for qPCR, related to STAR Methods**

**Captions for Videos S1 to S4**

Videos are provided as separate files.

**Video S1, related to Figure 5**

Cryo-electron tomogram and 3D tracing of nucleocapsids in a mumps viral factory in unstressed HeLa cells (scale bar: 100 nm, 6.74 nm-thick tomographic slices). Data were acquired with a Volta phase plate and defocus.

**Video S2, related to Figure 5**

Cryo-electron tomogram and 3D tracing of nucleocapsids in a mumps viral factory in HeLa cells at 1 h of 0.5 mM As(III) stress (scale bar: 100 nm, 6.74 nm-thick tomographic slice). Data were acquired with a Volta phase plate and defocus.

**Video S3, related to Figure 5**

Cryo-electron tomogram and 3D tracing of nucleocapsids in a mumps viral factory in HeLa cells at 6 h of 30 μM As(III) stress (scale bar: 100 nm, 6.52 nm-thick tomographic slice). Data were acquired with defocus only and radial filtering applied during IMOD tomogram reconstruction with the default values.

**Video S4, related to Figures 6 and 7**

EM maps and structural models showing different conformations of the authentic MuV nucleocapsid, *ex* and *in situ*.
