## Supplementary material for "Molecular mechanisms of stress-induced reactivation in mumps virus condensates": Key resources table

| REAGENT or RESOURCE | SOURCE | IDENTIFIER |
|---|---|---|
| Antibodies | | |
| Mouse monoclonal anti-MuV-N protein (clone 8H4) | Abcam | Cat#ab9876 |
| Goat anti-mouse IgG secondary antibody Alexa Fluor 488 (highly cross-adsorbed) | Invitrogen | Cat#A-11029 |
| Mouse monoclonal anti-GAPDH (clone 6C5) | Santa Cruz | Cat#sc-32233 |
| Goat anti-mouse IgG (H/L):HRP (multi species absorbed) | Bio-Rad | Cat#STAR117P |
| Bacterial and virus strains | | |
| Mumps virus Enders strain | Naturally persisting in the HeLa cell culture | NCBI: GU980052 |
| Chemicals, peptides, and recombinant proteins | | |
| Sodium arsenite | Sigma-Aldrich | S7400; CAS: 7784-46-5 |
| Potassium arsenate | Sigma-Aldrich | A6631; CAS: 7784-41-0 |
| 1,6-Hexanediol | Sigma-Aldrich | 240117; CAS: 629-11-8 |
| oligo $d(T)_{23}$ primer | New England Biolabs | Cat#S1327S |
| EDTA-free protease inhibitor cocktail | Roche | 04693132001 |
| PhosSTOP Phosphatase inhibitor cocktail | Roche | 4906845001 |
| Benzonase Nuclease, ultrapure | Millipore | E8263; CAS: 9025-65-4 |
| Ribonuclease inhibitor | Sigma-Aldrich | R1158 |
| Heparin sodium | Sigma-Aldrich | H3149; 9041-08-1 |
| Protein-A/Gold, EM-grade 10 nm | Electron Microscopy Sciences | Cat#50-281-97 |
| 1-µm crimson beads (FluoSpheres carboxylate-modified microspheres, 625/645) | Invitrogen | Cat#F8816 |
| Critical commercial assays | | |
| jetPRIME Transfection Reagent | Polyplus | Ref: 101000027 |
| RNeasy Micro Kit | Qiagen | Cat#74004 |
| ProtoScript II Reverse Transcriptase Kit | New England Biolabs | Cat#M0368 |
| TaqPath qPCR Master Mix, CG | Applied Biosystems | Cat#A15297 |
| RNA Nano 6000 Assay Kit | Agilent Technologies | Cat#5067-1511 |
| NEBNext Ultra II Directional RNA Library Prep Kit for Illumina | New England Biolabs | Cat#E7760 |
| SPRIselect beads | Beckman Coulter | B23318 |
| DNA High Sensitivity kit | Agilent Technologies | 5067-4626 |
| Qubit dsDNA High Sensitivity kit | Invitrogen | Cat#Q32851 |
| Bradford Protein Assay Dye | Bio-Rad | Cat#5000006 |
| Deposited data | | |
| PIV5 nucleocapsid-RNA complex | (Alayyoubi et al., 2015) | PDB: 4XJN |
| PIV5 L-P complex | (Abdella et al., 2020) | PDB: 6V85 |
| EM map of PIV5 L-P complex | (Abdella et al., 2020) | EMD-21095 |
| Cryo-ET of viral factory at non-stressed condition | This study | EMD-13165 |
| Cryo-ET of Viral factory at 1 h of 0.5mM As(III) stress | This study | EMD-13166 |

| | | |
|---|---|---|
| Cryo-ET of Viral factory at 6 h of 30 μM As(III) stress | This study | EMD-13167 |
| EM map for isolated nucleocapsids, majority class | This study | EMD-13133 |
| EM map for isolated nucleocapsids, minority class | This study | EMD-13136 |
| EM map for straight cellular nucleocapsids | This study | EMD-13137 |
| Model for isolated nucleocapsids, majority class | This study | PDB: 7OZR |
| Raw micrographs for isolated nucleocapsids | This study | EMPIAR-10751 |
| Mass spectrometry data | This study | ProteomeXchange: PXD026799 |
| Reference proteome sequence for *Homo sapien* | Uniprot | UP000005640 |
| Reference genome NCBI | (Young et al., 2009) | GU980052 |
| **Experimental models: Cell lines** | | |
| HeLa: G3BP1-mCherry BAC in HeLa Kyoto cell line | (Guillen-Boixet et al., 2020) | MCB_ky_7510 (C-terminal mCherry tag; FACS sorted) |
| HeLa-MuV: G3BP1-mCherry BAC in HeLa Kyoto cell line persistently infected by mumps virus | Parental cell line from (Guillen-Boixet et al., 2020) | MCB_ky_7510 (C-terminal mCherry tag; FACS sorted) |
| **Recombinant DNA** | | |
| pcDNA3.1-MuV-P | This study (synthesized by GeneArt/Life Technologies) | N/A |
| pcDNA3.1-MuV-P-EGFP | This study | N/A |
| pcDNA3.1-MuV-N | This study (synthesized by GeneArt/Life Technologies) | N/A |
| **Software and algorithms** | | |
| SerialEM | (Mastronarde, 2005) | https://bio3d.colorado.edu/SerialEM/ |
| Dose-symmetric tomography acquisition scheme | (Hagen et al., 2017) | N/A |
| IMOD package | (Kremer et al., 1996) | https://bio3d.colorado.edu/imod/ |
| 3DCT 2.2.2 | (Arnold et al., 2016) | https://3dct.semper.space/ |
| Amira 6.7 and 2019.4 | ThermoFisher Scientific | https://www.fei.com/software/amira-release-notes/ |
| Warp 1.0.9 | (Tegunov and Cramer, 2019) | http://www.warpem.com |
| M 1.0.9 | (Tegunov et al., 2021) | http://www.warpem.com |
| Matlab R2019a | MathWorks | https://www.mathworks.com |
| Dynamo 1.1.401 | (Castano-Diez et al., 2012) | https://wiki.dynamo.biozentrum.unibas.ch |
| TOM | (Nickell et al., 2005) | N/A |
| AV3 | (Forster and Hegerl, 2007) | N/A |
| RELION 3.0 and 3.1 | (Zivanov et al., 2018) | https://github.com/3dem/relion |

| | | |
|---|---|---|
| I-TASSER | (Roy et al., 2010) | https://zhanglab.dcmb.med.umich.edu/I-TASSER/ |
| Coot 0.9 | (Emsley et al., 2010) | https://www2.mrc-lmb.cam.ac.uk/personal/pemsley/coot/ |
| Phenix 1.18-3845 | (Liebschner et al., 2019) | https://phenix-online.org |
| Chimera 1.13.1 | (Pettersen et al., 2004) | https://www.cgl.ucsf.edu/chimera/ |
| ChimeraX 1.1.1 | (Pettersen et al., 2021) | https://www.cgl.ucsf.edu/chimerax/ |
| Fiji 2.1.0 | (Schindelin et al., 2012) | https://imagej.net/software/fiji/ |
| Imaris 9.5.1 | Oxford Instruments Group | https://imaris.oxinst.com |
| Image Lab 6.1 | Bio-Rad | https://www.bio-rad.com |
| R 3.6.1 | R | https://www.r-project.org |
| RStudio 1.4.1103 | RStudio | https://www.rstudio.com |
| Prism 6.0c | GraphPad | https://www.graphpad.com |
| IUPred2A | (Meszaros et al., 2018) | https://iupred2a.elte.hu/ |
| PSIPRED | (Jones, 1999) | http://bioinf.cs.ucl.ac.uk/psipred/ |
| BWA-MEM 0.7.17-r1188 | N/A | https://github.com/lh3/bwa |
| Picard tool 2.9.0 | Broad Institute of MIT and Harvard | https://broadinstitute.github.io/picard |
| FreeBayes 1.1.0-3 | N/A | https://github.com/freebayes/freebayes |
| isobarQuant | (Breitwieser et al., 2011) | https://github.com/protcode/isob |
| Mascot 2.4 | Matrix Science | https://www.matrixscience.com |
| vsn | (Huber et al., 2002) | https://bioconductor.org/packages/release/bioc/html/vsn.html |
| limma | (Ritchie et al., 2015) | https://bioconductor.org/packages/3.14/bioc/html/limma.html |
| clusterProfiler | (Yu et al., 2012) | https://bioconductor.org/packages/3.14/bioc/html/clusterProfiler.html |
| 3D-Unet | N/A | https://github.com/irenedet/3d-unet/tree/7bc343971bdb818c5de90570b83731c8d77cde04 |
